# Structural brain development: a review of methodological approaches and best practices

**DOI:** 10.1101/153718

**Authors:** Nandita Vijayakumar, Kathryn L. Mills, Aaron Alexander-Bloch, Christian K. Tamnes, Sarah Whittle

**Affiliations:** Department of Psychology, University of Oregon, Eugene, USA; Department of Psychiatry, Yale University School of Medicine, New Haven, Connecticut, USA; Department of Psychology, University of Oslo, Oslo, Norway; Melbourne Neuropsychiatry Centre, Department of Psychiatry, The University of Melbourne and Melbourne Health, Melbourne, Australia; Melbourne School of Psychological Sciences, The University of Melbourne, Melbourne, Australia

**Keywords:** structural MRI, brain development, longitudinal analyses, methodology

## Abstract

Continued advances in neuroimaging technologies and statistical modelling capabilities have improved our knowledge of structural brain development in children and adolescents. While this has provided an increasingly nuanced understanding of brain development, the field is still plagued by inconsistent findings. This review highlights the methodological diversity in existing longitudinal magnetic resonance imaging (MRI) studies on structural brain development during childhood and adolescence, and addresses how such variation might contribute to inconsistencies in the literature. We discuss the impact of method choices at multiple decision points across the research process, from study design and sample selection, to image processing and statistical analysis. We also highlight the extent to which different methodological considerations have been empirically examined, drawing attention to specific areas that would benefit from future investigation. Where appropriate, we recommend certain best practices that would be beneficial for the field to adopt, including greater completeness and transparency in reporting methods, in order to ultimately develop an accurate and detailed understanding of normative child and adolescent brain development.

## 1. Introduction

Over the past two decades we have learnt a great deal about normative structural brain development during childhood and adolescence with the application of magnetic resonance imaging (MRI) in longitudinal projects. While the pioneer studies published in the 1990s and 2000s continue to be among the most influential and often cited, more recent investigations have provided complementary, but also sometimes contradictory findings on normative structural brain development. This paper aims to highlight potential methodological causes of inconsistencies in findings on structural brain development across studies, focusing on the impact of specific method choices at multiple decision points along the research process, from study design and sample selection, to image processing and statistical analysis.

A growing number of longitudinal projects aim to characterize typical structural brain development in children and/or adolescents, many of which are summarized in Table 1. While some characteristics of these projects overlap, differences are also evident for instance in sample size, age range, number of repeat assessments, and study design. Multiple studies commonly arise from each dataset, which often differ in methodology, as outlined in Table 2. The diversity of MRI processing techniques, structural measures of interest and statistical analytic methods used across these studies is a demonstration of the productivity and ever-evolving nature of the fields of neuroimaging and developmental neuroscience. However, it is also important to consider how different methods impact the results of studies investigating typical brain developmental trajectories. Following a brief overview of current findings, we explore how each methodological step, from study design and image acquisition to model fitting, might influence findings and conclusions. We focus specifically on longitudinal studies of typically developing children (5 years and older) and adolescents. Younger age ranges were excluded due to methodological issues that are either unique or exemplified in this population (e.g., techniques to reduce anxiety and movement, such as scanning during natural sleep, the availability of child-appropriate equipment, and use of appropriate analytic techniques such as pediatric brain templates; as described by Raschle et al., 2012. Further details regarding the search strategy and inclusionary criteria is presented in Box 1.

#### Box 1

We searched PubMed using the following terms: brain AND development AND (childhood OR adolescence) AND (structure OR thickness OR volume OR surface area OR gyrification) AND MRI, to identify studies published in this field to date (January 2017). Inclusionary criteria for the review were: i) sample age range predominantly encompassing mid-childhood (5 years) to young adulthood, ii) focused on normative development, iii) use of structural MRI to examine grey matter brain structure, iv) longitudinal study design, v) total number of scans greater than 50, and vi) written in English. The reference lists of identified articles were also searched for further relevant articles. Identified studies are summarized in Table 2.

**Table 1.** Overview of longitudinal structural MRI datasets.

| **Project** | **Age-range, years** | **n participants (longitudinal<sup>a</sup>)** | **N scans** | **Average scans per participant (i.e. N/n)** | **Range scans** | **Longitudinal study design (ALD vs SCD)** | **Field strength / voxel size** |
| --- | --- | --- | --- | --- | --- | --- | --- |
| **Alberta Canada sample** | 5 - 32 | 103 (103) | 221 | 2.15 | 1 - 4 | ALD | 1.5T<br>1x1x1 |
| **BrainSCALE UMCU - NTR** | 9 - 13 | 224 (178) | 346 | 1.54 | 1 - 3 | SCD | 1.5T<br>1x1x1.2 |
| **Braintime** | 8 - 28 | 271 (241) | 680 | 2.51 | 1 - 3 | ALD | 3T<br>0.875x0.875x1.2 |
| **Leonard Florida sample** | 5 - 11 | 45 (45) | 90 | 2.00 | 2 | ALD | 1.5T<br>0.98x0.98x1.25 |
| **Mother-Child Cohort Study** | 4 - 10 | 428 (304) | 732 | 1.71 | 1 - 2 | ALD | 1.5T<br>1.25x1.25x1.2 |
| **Neurocognitive Development** | 8 - 25 | 191 (148) | 407 | 2.13 | 1 - 3 | ALD | 1.5T<br>1.25x1.25x1.2 |
| **NICHE cohort** | 7 - 23 | 147 (53) | 233 | 1.59 | 1 - 3 | ALD | Two scanners: 1.5T<br>1x1x1.2 |
| **NIH MRI Study of Normal Brain Development** | 5 - 22 | 538 (527) | 1381 | 2.56 | 1 - 3 | ALD | 6 scanners: all 1.5T<br>In-plane 1x1, slice thickness ranged from 1-1.8mm |
| **NIMH Child Psychiatry Branch** | 3 - 30 | 647 (376) | 1274 | 1.93 | 1 - 7 | ALD | 1.5T<br>0.94x0.94x1.5 |
| **Orygen Adolescent Development Study** | 11 - 20 | 166 (128) | 367 | 2.21 | 1 - 3 | SCD | 2 scanners: both 3T<br>1: 0.48x0.48x1.5<br>2: 0.9x0.9x0.9 |
| **University of Minnesota cohort** | 9 - 24 | 149 (149) | 298 | 2.00 | 2 | ALD | 3T<br>1x1x1 |
| **University of Pittsburgh cohort** | 10 - 14 | 126 (81) | 226 | 1.79 | 1 - 2 | SCD | 3T<br>1x1x1 |

NB: This table only reports longitudinal datasets that have been published, including both projects that are completed and still ongoing. Details were acquired by contacting investigators, or from studies published using the datasets. <sup>a</sup> Longitudinal participants refers to the number of participants that have 2 or more scans. ALD = Accelerated longitudinal; SCD = Single cohort design; NIH = National Institute of Health; NIMH = National Institute of Mental Health; UMCU – NTR = University Medical Center in Utrecht - Netherlands Twin Register

**Table 2.** Details of longitudinal studies investigating normative structural brain development between childhood and young adulthood.

| Study (Project) | N (males) | N Scans, n per subject, approximate interval | Age (y) | Image processing software (version) | Measures: vol/sa/ct/others | Specificity of analyses | Index of analyses: absolute or change values, whole brain correction | Statistical analyses: analysis method (software), effects, model fit, trajectories, multiple comparison |
| --- | --- | --- | --- | --- | --- | --- | --- | --- |
| Lebel & Beaulieu, 2011 (Alberta, Canada) | 103 (51) | 221, 2-4 per subject, 4 year | 5 - 32 | FreeSurfer | Volume | Global | Absolute values and difference score for within subject change (based on change >1SD) | Mixed models<br>Effects: age, controlling for sex<br>Model selection: step-down<br>Trajectories: linear, quadratic |
| Zhou et al., 2015 (Albeta, Canada) | 90 (42) | 180, 2 per subject, 4 year | 5 - 32 | Civet 1.1.11 | CT, SA | Global and lobar | Absolute values and difference score for within subject change (based on change >1SD) | Student's t-test compared mean thinning rates across age groups<br>Kruskal–Wallis test of differences in ratio of increased/decrease/no change between age groups<br>Trajectories: linear |
| Swagerman et al., 2014 (BRAINSscale) | 224 (112) | 346, 1-2 per subject, 3 year | 9 - 12 | FreeSurfer 5.1 | Volume | Segmentation | Absolute values and ICV-corrected | Bivariate analyses of twin data: OpenMX<br>Effects: age within each sex, sex at each time point<br>MC: Bonferroni for number of independent dimensions in data |
| van Soelen et al., 2012 (BRAINSscale) | 113 (60) | 226, 2 per subject, 3 year | 9 - 13 | Automated: Peper et al., 2008; Brouwer et al., 2010. CLASP algorithm | CT | Vertex-wise | Change (difference) | One sample t-test<br>Effects: sex, controlling for handedness and duration of scan-interval.<br>Trajectories: linear<br>MC: FDR |
| Aleman-Gomez et al., 2013 (Child and adolescent first-episode psychosis study) | 52 (32) | 104, 2 per subject, 2 year | 11 - 17 | FreeSurfer 5.1 LP; BrainVisa 4.2.1 | CT, SA, GI, gyral WM thickness, convex hull SA, sulcal length/depth/width. | Global and lobar | Percentage change (average or summed across hemispheres) | GLM: SPSS<br>One sample t-test<br>Effects: lobe, age, sex, interaction of age and sex, scanner, time between acquisitions.<br>MC: FDR |
| Sowell et al., 2004<br>(Leonard Florida) | 45 (23) | 90,<br>2 per subject,<br>2 year | 5 - 11 | Automated:<br>MacDonald et al.,<br>1994, Thompson et al.,<br>2000a, Sowell et al.,<br>2001b;<br>Manual tracing of<br>sulcal delineation | CT and brain<br>growth<br>(distance from<br>center of<br>brain/hemispher<br>e) | Vertex-wise,<br>lobar, and<br>perisylvian ROI | Change (i.e.<br>difference) | One-sample t-test<br>MC: permutation testing |
| Tamnes et al.,<br>2013 (NCD) | 85 (47) | 170,<br>2 per subject,<br>2.6 year | 8 -22 | FreeSurfer 5.1;<br>QUARC | Volume | Vertex-wise and<br>segmentation | Percentage change<br>(vertex and<br>subcortical) | GLM (change differs from zero): FreeSurfer,<br>SPSS, R<br>Effects: age, sex, and interaction, controlling<br>for scan interval<br>Trajectories: linear<br>Assumption-free models used for description<br>(no statistical testing)<br>MC: FDR & Bonferroni |
| Wierenga et al.,<br>2014a (NICHE) | 135 (92) | 201,<br>1 - $\geq 3$ per<br>subject,<br>1.5-5.5 year | 7 - 23 | FreeSurfer 5.1 | CT, SA, CV | Parcellation | Absolute | Mixed models<br>Effects: Age, sex, and interactions<br>Model selection: Step-down for age, BIC for<br>sex<br>Trajectories: linear, quadratic, cubic |
| Wierenga et al.,<br>2014b (NICHE) | 147 (94) | 223<br>$\geq 1$ per subject,<br>1.5-5.6 year | 7 - 23 | FreeSurfer 5.1 | Volume | Segmentation | Absolute | Mixed models<br>Effects: age, sex, and interactions<br>Model selection: stepdown for age, BIC for<br>sex<br>Trajectories: linear, quadratic, cubic |
| Aubert-Broche et<br>al., 2013 (NIH<br>MRI) | 292 | 882,<br>2-4 per<br>subject,<br>2 year | 4 - 19 | Longitudinal pipeline<br>("LL method") | Volume | Global and<br>regional/segmenta<br>tion | Absolute | Mixed models: R<br>Effects: age, sex, and interactions<br>Model selection: AIC<br>Trajectories: linear, quadratic |
| Cao et al., 2015<br>(NIH MRI) | 303 (142) | 418,<br>1-2 per<br>subject,<br>2 year | 5 - 18 | FreeSurfer | Volume | Parcellation and<br>segmentation | Absolute | LASSO: multivariate linear regression<br>Effects: age |
| Ducharme et al.,<br>2015 (NIH MRI) | 384 (343) | 753,<br>1-3 per<br>subject,<br>2 year | 5 - 22 | CIVET 1.1.11 | SA, CV | Vertex-wise and<br>lobar | Absolute | Mixed models: SurfStat, R<br>Effects: age with and without controlling for<br>WBV<br>Model selection: Step-down (vertex) & AIC<br>(lobar)<br>Trajectories: linear, quadratic, cubic |
| Ducharme et al., 2016 (NIH MRI) | 383 (343) | 753,<br>1-3 per subject,<br>2 year | 5 - 22 | CIVET 1.1.11 | CT | Vertex-wise and lobar | Absolute | Mixed models: SurfStat, R<br>Effects: age, sex, with and without controlling for WBV<br>Model selection: Step-down (vertex) & AIC (lobar)<br>Trajectories: linear, quadratic, cubic |
| Krongold et al., 2015 (NIH MRI) | 335 (155) | 724,<br>1-2 per subject | 4 - 22 | FreeSurfer 5.3 (LP) | CT, SA, CV | Parcellation | Absolute | Mixed models: R (lme4)<br>Effects: age with sex as nuisance regressor<br>Trajectories: linear |
| Nie et al., 2013 (NIH MRI) | 445 (127) | 951 | 3 - 20 | Automated: Zhang et al. 2001 & Liu et al. 2008 | CT | Regional | Absolute | Linear regression |
| Giedd et al., 1999 (NIMH CPB) | 145 (89) | 280 scans,<br>1-5 per subject,<br>2 year | 4 - 22 | Automated: Zijdenbos et al., 2002 | GM Volume | Lobar | Absolute | Mixed models<br>Effects: age, sex and interactions<br>Model selection: Step-down<br>Trajectories: linear, quadratic |
| Gogtay et al., 2004 (NIMH CPB) | 13 (6) | 52,<br>≥3 per subject,<br>2 year | 4 - 21 | Automated: Thompson et al. 2001 | GM volume, GM density | Lobar and vertex-wise | Absolute | Mixed models<br>Effects: age<br>Model selection procedure: Step-down<br>Trajectories: cubic, quadratic, linear |
| Gogtay et al., 2006 (NIMH CPB) | 31 (16) | 100<br>≥2 per subject,<br>2 year | 4 - 25 | Manual tracing from single individual; surface mesh applied to hippocampus | GM Volume | ROI and vertex-wise | Absolute | Mixed models<br>Effects: age, sex and interactions; WBV used as a covariate<br>Model selection procedure: Step-down<br>Trajectories: cubic, quadratic, linear |
| Harezlak et al., 2005 (NIMH CPB) | 300 (159) | 619,<br>1-5 per subject | 3 - 25 |  | Volume | Total cerebral volume and ROIs | Absolute | Parametric (polynomial) vs. semiparametric (reduced rank penalized regression models)<br>Effects: age, sex |
| Lenroot et al., 2007 (NIMH CPB) | 387 (209) | 829<br>1-7 per subject,<br>2 year | 3 - 27 | Automated Nonlinear Image Matching and Anatomical Labelling | GM volume, WM volume | Global and lobar | Absolute and percentage change | Mixed models<br>Effects: sex, with and without adjustment for WBV at the same age<br>Model selection: Step-down<br>Trajectories: cubic, quadratic and linear |
| Mills et al., 2014a (NIMH CPB) | 33 (23) | 152,<br>3-6 per subject,<br>2 year | 7 - 30 | Freesurfer 5.3 (LP) | Volume | ROI | Absolute | Mixed models: R<br>Effects: age, and interactions<br>Model selection: AIC<br>Trajectories: linear, quadratic, cubic |
| Mills et al., 2014b | 288 (164) | 857 (ATC: | 7 - 30 | Freesurfer 5.1 | CT, SA, CV | ROI | Absolute | Mixed models: R |
| (NIMH CPB) | ATC: 221 | 447,<br>2-7 per<br>subject,<br>2 year |  |  |  |  |  | Effects: age, sex, and interactions<br>Model selection: AIC<br>Trajectories: linear, quadratic, cubic |
| Raznahan et al.,<br>2011a (NIMH<br>CPB) | 647 (328) | 1274,<br>1 - $\geq 3$ per<br>subject,<br>2 year | 3 - 30 | MNI anatomical<br>pipeline | CT, SA, CV,<br>GI, CHA | Global | Absolute values and<br>rate of change | Mixed models: R<br>Effects: age, sex, and interactions<br>Model selection: Step-down for age,<br>likelihood ratio tests for sex<br>Trajectories: linear, quadratic, cubic |
| Raznahan et al.,<br>2014 (NIMH<br>CPB) | 618 (312) | 1171,<br>1 - $\geq 3$ per<br>subject,<br>2 year | 5 - 25 | Volume: MAGeT<br>Brain<br>SA: Marching cubes<br>and AMIRA<br>CV: CIVET | Volume, SA | Segmentation and<br>global CV | Absolute | Mixed models: R<br>Effects: age, sex, and interactions<br>Model selection: Step-down for age,<br>likelihood ratio tests for sex<br>Trajectories: linear, quadratic, cubic |
| Shaw et al., 2008<br>(NIMH CPB) | 375 (196) | 764,<br>1-$\geq 4$ per<br>subject,<br>2 year | 3 - 33 | Automated: Zijdenbos<br>et al., 2002 | CT | ROI and<br>vertexwise | Absolute | Mixed models<br>Effects: age<br>Model selection: Step down<br>Trajectories: cubic, quadratic and linear |
| Tiemeier et al.,<br>2010 (NIMH<br>CPB) | 50 (25) | 183,<br>$\geq 3$ per subject,<br>2 year | 5 - 24 | Automated: Zijdenbos<br>et al., 2002;<br>Manual tracing of<br>subregions | Volume | Parcellation of<br>cerebellum | Absolute | Mixed models<br>Effects: sex, with and without adjustment for<br>WBV<br>Model selection: Step-down<br>Trajectories: linear, quadratic, cubic |
| Dennison et al.,<br>2013 (OADS) | 60 (32) | 120,<br>2 per subject,<br>4 year | 11 - 18 | FreeSurfer 5.1 | Volume | Segmentation | Absolute values and<br>WBV-corrected | Hierarchical linear models: Stata<br>Effects: Age, hemisphere, sex, and<br>interactions<br>Trajectories: linear<br>MC: B-Y method |
| Vijayakumar et<br>al., 2016 (OADS) | 90 (49) | 192,<br>1 - 3 per<br>subject,<br>3 year | 11 - 20 | FreeSurfer 5.3 (LP) | CT, SA, CV | Parcellation and<br>vertex-wise | Absolute | Mixed models: SPSS, FreeSurfer LMM<br>toolbox<br>Effects: Age, sex, and interactions<br>Model selection: BIC (parcellation), step-down (vertex)<br>Trajectories: linear, quadratic<br>MC: FDR |
| Sullivan et al.,<br>2011 (Stanford<br>Research Institute) | 28 (16) | 56,<br>2 per subject,<br>7.3 months | 11 - 14 | FSL FAST | Volume | Lobar and ROI | Absolute | Percent change<br>Effects: age, sex |
| Mutlu et al., 2013<br>(Switzerland) | 137 (68) | 209,<br>1-4 per<br>subject | 6 - 30 | FreeSurfer | CT, GI | Vertex-wise | Absolute | Mixed models: Matlab (nlmefit)<br>Effects: age, sex, and interactions<br>Model selection: BIC for age, LRT for sex<br>Trajectories: linear, quadratic and cubic<br>MC: Monte Carlo simulation in FreeSurfer |
| Tanaka et al.,<br>2012 (Toyama,<br>Japan) | 114 (60) | 209,<br>1-4 per<br>subject | 1m - 25 | Manual tracing | Volume | Global and lobar | Absolute values and<br>ICV-corrected | Linear regression<br>Effects: age, controlling for sex and<br>hemisphere<br>Model selection: R squared<br>Trajectories: linear, quadratic and cubic |
| Urosevic et al.,<br>2012 (University<br>of Minnesota) | 149 | 298,<br>2 per subject,<br>2 year | 9 - 26 | FreeSurfer 4.5 (LP) | Volume | ROIs | WBV-corrected | Repeated-measures ANCOVAs: SPSS<br>Effects: time, age (covariate), sex, time*age,<br>time*sex, controlling for scanner upgrade<br>Trajectories: linear |
| Fjell et al., 2015<br>(NCD, MCCN<br>Study, Cognitive<br>and Plasticity<br>through the<br>Lifespan) | 974 (466) | 1633,<br>1-3 per<br>subject,<br>2.5 year | 4 - 89 | FreeSurfer 5.3 (LP) | CT | Parcellation based<br>on genetic<br>clustering | Absolute and<br>percentage change | General additive mixed models: R; Linear<br>mixed models: Matlab<br>Effects: age (sex not found to influence<br>preliminary results)<br>Model fit: AIC and BIC<br>Trajectories: linear, smoothing spline<br>MC: FDR |
| Mills et al., 2016<br>(Braintime, NIMH<br>CPB, NCD,<br>Pittsburgh) | 391 (191) | 852<br>≥2 per subject | 7 - 30 | FreeSurfer 5.3 (LP) | Volume | Global | Absolute | Mixed models: R<br>Effects: age, sex, with and without controlling<br>for ICV or WBV<br>Model selection: AIC<br>Trajectories: linear, quadratic, cubic |

NB: Inclusionary criteria are presented in Box 1. Studies are grouped by project, and subsequently ordered by author surname and year published. AIC = Akaike Information Criteria; ANCOVA = analysis of covariance; BIC = Bayesian information criterion; B-Y = Benjamini-Yekutieli; CT = cortical thickness; CV = cortical volume; FDR = false discovery rate; GI = gyrification index; GLM = general linear model; GM = grey matter; ICV = intracranial volume; LMM = linear mixed models; LP = longitudinal processing; LRT = likelihood ratio test; MCCN = Mother Child Cohort Study; NCD = Neurocognitive Development; NIH = National Institute of Health; NIMH CPB = National Institute of Mental Health Child Psychiatry Branch; OADS = Orygen Adolescent Development Study; ROI = region of interest; SA = surface area; WBV = whole brain volume; WM = white matter

## 2. Overview of findings

Initial studies from the National Institute of Mental Health Child Psychiatry Branch (NIMH CPB) described inverted-U-shaped growth trajectories of cortical volumetric grey matter development (Giedd et al., 1999; Lenroot et al., 2007), reporting peak volumes around early adolescence that distinguished periods of growth during childhood from reductions during adolescence. However, results from subsequent studies using other longitudinal datasets have not identified such “peaks”; many studies report continued reductions in grey matter volumes from late childhood into adolescence (Aubert-Broche et al., 2013; Tamnes et al., 2013; Wierenga et al., 2014b). Studies have also reported temporal patterns of maturation, including rostral-to-caudal waves of growth in the corpus callosum (Thompson et al., 2000) and posterior-to-anterior growth in the frontal lobe (Gogtay et al., 2004), although few have attempted to replicate these effects in different samples. In contrast to cortical grey matter volume, studies have consistently reported an increase in white matter volume across childhood and adolescence (Aubert-Broche et al., 2013; Lebel and Beaulieu, 2011; Mills et al., 2016). A recent study highlighted convergence in developmental patterns of grey and white matter volume across four longitudinal studies when employing the same preprocessing stream and analytic methods (see Figure 1; Mills et al., 2016). However, others have noted variability in developmental trajectories at higher anatomical resolutions such as the vertex level (Mutlu et al., 2013; Vijayakumar et al., 2016).

**Figure 1.**
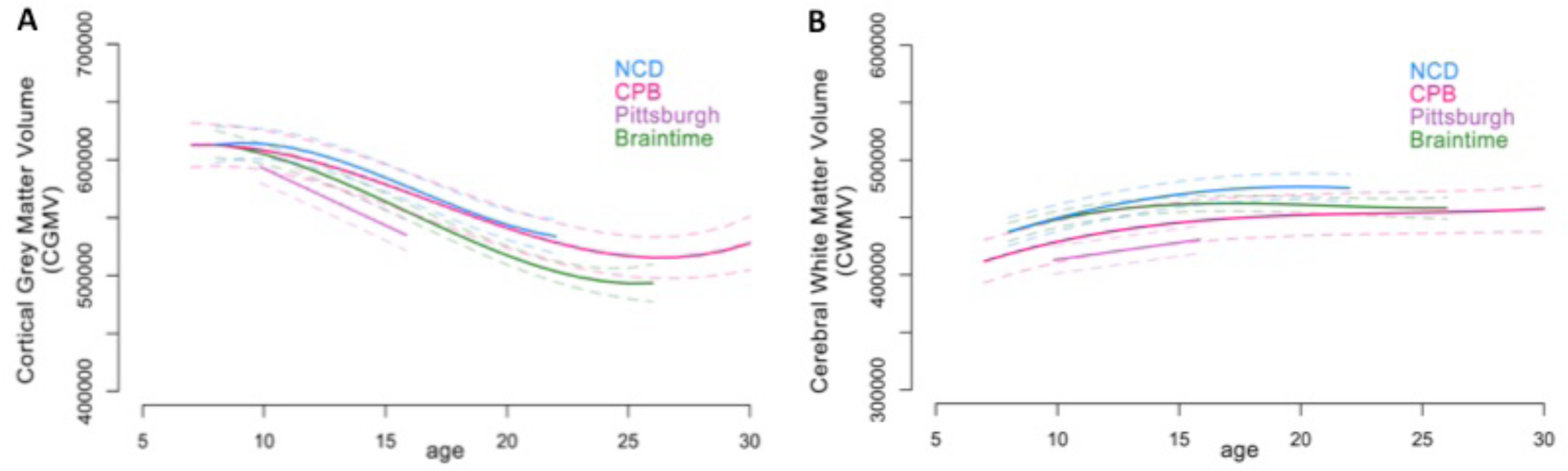
Development of a) cortical grey matter volume and b) cortical white matter volume across four longitudinal datasets. NCD = Neurocognitive Development, CPB = (National Institute of Health) Child Psychiatry Branch. Adapted from Mills et al. (2016).

Over time, there has been an increasing emphasis on the examination of the subcomponents of cortical volume: *thickness*, defined as the distance between the white matter/grey matter cortical boundary and grey matter/CSF cortical boundary; and *surface area*, defined as the area of one of these two boundaries (or surfaces). While the majority of studies have identified reductions in cortical thickness between childhood and adulthood (e.g., Wierenga et al., 2014b), some have found nonlinear global development (e.g., Raznahan et al., 2011b). In contrast, studies consistently report global surface area increasing between childhood and early adolescence (Raznahan et al., 2011b; Wierenga et al., 2014b) before decreasing across the rest of the second decade (Alemán-Gómez et al., 2013). Regional differences have also been reported for these subcomponents of cortical volume (Mutlu et al., 2013; Tamnes et al., 2017; Vijayakumar et al., 2016; Wierenga et al., 2014a).

A small number of studies have investigated measures of gyri and sulci structure. The exposed outer cortical surface area, referred to as the convex hull area (CHA), has been found to show both quadratic (Raznahan et al., 2011b) and linear (Alemán-Gómez et al., 2013) reductions with age, while linear reductions in the degree of gyrification have been found more consistently (ratio of total cortical surface area to CHA: gyrification index (GI); Alemán-Gómez et al., 2013; Raznahan et al., 2011b). However, one vertex-wise investigation reported that GI might not change in certain parts of the medial surface between childhood and adolescence (Mutlu et al., 2013).

As studies try to unpack the complex relationships between these different brain measures, we are gaining a more nuanced understanding of how brain structure develops. While there is, overall, convergence in findings on a broad scale (i.e., overall direction of change), inconsistencies are evident when considering details such as the precise shape of developmental trajectories, presence/location of peaks, regional variability and sex differences. Following, we discuss each of the likely major methodological contributions to these inconsistences.

## 3. Study and sampling design

### 3.1 Study design

Since maturation (i.e., age) cannot be randomly assigned to participants in studies investigating brain development, it represents a correlational or quasi-independent variable. The resultant quasi-experimental research designs can be broadly grouped into one of three categories: cross-sectional, complete longitudinal or single cohort design (SCD), and accelerated longitudinal design (ALD; Appelbaum and McCall, 1983; Bordens and Abbott, 2013). A limitation common to these designs is that a causal relationship cannot be directly inferred between age and the variables of interest, as third (confounding) variables cannot be fully accounted for.

Inferences about developmental processes from studies with cross-sectional designs, where different participants at different ages are compared, can be misleading (Kraemer et al., 2000). Also, because of large individual differences in brain structure, longitudinal designs with repeated measurements of the same participants have greatly increased statistical power (Steen et al., 2007). Therefore, cross-sectional studies are not reviewed here. SCD studies, where all participants begin at the same age and are followed across the entire age-range of interest, have the advantage of simplicity and are more amenable to certain modelling techniques (King et al., this issue), but are time-consuming, costly, and may not be feasible for broad age ranges. Of the 34 studies reviewed (Table 2), only 4 were SCD studies: two studies from the same project focus on a narrow age-range (9-13 years; Swagerman et al., 2014; van Soelen et al., 2012), and a further two studies from the same project focus on a broader age range (11-18 and 11-20; Dennison et al., 2013; Vijayakumar et al., 2016).

Because of the limitations of SCD studies, nearly all longitudinal studies of structural brain development in childhood and adolescence have used ALD. In ALD, participants begin at different ages or years and contribute data to only part of the age-range of interest. These designs thus include both a cross-sectional and a longitudinal component. Compared to SCD, ALD can cover the age-range of interest with a shorter study duration, they are less affected by participant dropout (attrition), and this dropout tends to be less systematically related to age. ALD is also less vulnerable to the effects of unforeseen method or procedure changes during the data collection period (e.g., scanner change or upgrades); these confounding variables in SCD are often more systematically related to age. SCD also confounds age with potential cohort effects. However, the major trade-off of ALD is the inherent missing data for each participant (Galbraith et al., 2017), and some individuals may only contribute a single (i.e., cross-sectional) data point to the study.

ALD studies differ widely in the number of participants and measurements, and the frequency and timing of measurements, factors that have implications for the duration and cost of the study, and also the statistical analyses (Galbraith et al., 2017). Many ALD developmental imaging studies appear to be structured such that individuals enter the study at pre-selected ages (i.e., age cohorts), which together span the age range of interest. The spans of the age cohorts overlap, and subjects are followed longitudinally over a shorter time span relative to the entire age range (Bell, 1954). However, it is of note that studies rarely describe this information in detail, and whether the design was tailored for separating the effects of age, cohort and/or time of measurement (see Appelbaum and McCall, 1983). Critically, small samples not only reduce the chance of detecting a true effect, but also reduce the likelihood that a statistically significant result reflects a true effect. The consequences of this are unreliable research and overestimated effect sizes (Button et al., 2013). In the current review, we have not included small longitudinal studies defined as those analyzing fewer than 50 scans.

### 3.2 Sample size and scan numbers

The sample sizes of the 34 studies included in Table 2 were highly variable. The number of participants ranged from 13 to 974, and the number of scans from 52 to 1633 (note that the largest study also included adults). Mean number of scans per participant ranged from 1.3 to 4.0, but to our knowledge, only 16 of 34 studies had on average more than two scans per participant, and only 3 (Gogtay et al., 2004; Mills et al., 2014a; Tiemeier et al., 2010) had three or more scans per participant on average. Thus, although several studies include relatively large samples, the amount of longitudinal data is generally low compared to many other areas of research, especially when considering that all except five of the ALD studies focused on an age-range of 13 or more years, with scan intervals typically being only a few years or less.

### 3.3 Sample characteristics

A number of important questions must also be addressed when choosing and recruiting participants (Bordens and Abbott, 2013), such as deciding upon the target population and the sampling and recruitment procedures, and defining eligibility and exclusionary criteria (Greene et al., 2016). As imaging studies of typical brain development rely upon volunteers that are willing to undergo MRI scans multiple times, they typically include non-random samples from subpopulations of the actual target population. Samples are usually relatively socioeconomically advantaged, have relatively high IQ, and are comprised of mostly Caucasian participants. As one exception, the NIH MRI Study of Brain Development used a population-based sampling method to ensure their sample was socio-demographically representative of the population.

Currently, the lack of detailed characterizations and reporting of the sampling procedure and final sample (e.g., approximately 40% of reviewed studies did not report sample IQ) prevents a good understanding of the generalizability of findings to the population. Further, oftentimes, studies arising from the same dataset use different subsamples and descriptions of study-specific selection criteria are not clear.

## 4. Image acquisition strategies and parameters

### 4.1 Strategies

Before performing MRI of children/adolescents, it is essential to systematically prepare the participant using, for instance, age-appropriate instructional videos or mock-scan. During the scan, multiple strategies can be implemented to create a good experience and obtain adequate data based on the need of the participant, such as having a parent present in the scanner room, playing a movie of their choice, and talking to them via the intercom between sequences (Greene et al., 2016). Optimization of the physical environment, for example with head cushions, may also increase subject comfort and decrease in-scanner motion.

Motion-related artefact can, to some extent, be mitigated by image acquisition methods. Perhaps most importantly, simple reductions in scanning time increase the probability of children remaining still throughout a scan. More involved methods can be broadly divided into retrospective techniques based on computational processing of scans (e.g., Atkinson et al., 1999) and prospective techniques that actually modify pulse sequences in response to detected motion (e.g., White et al., 2010). Even without explicit correction, tracking in-scanner motion, either via MR technology (Korin et al., 1990) or independently using external sensors (Qin et al., 2009), provides important information that can potentially be used in subsequent quality control (QC; see section 5 below).

### 4.2 Acquisition parameters

Most of the reviewed studies used 1.5T scanners, while more recently started projects typically use 3T scanners (i.e., only four of the 33 studies listed in Table 2 were performed only using 3T scanners (Dennison et al., 2013; Sullivan et al., 2011; Urosevic et al., 2012; Vijayakumar et al., 2016)). In addition, two studies included scans from both 1.5T and 3T, and analyzed them either independently (Mills et al., 2016) or together (Mutlu et al., 2013). Generally, higher field strength gives higher signal to noise ratio and improved spatial resolution at a fixed scan time, but some artefacts also become more prominent (Bernstein et al., 2006; Tijssen et al., 2009).

All studies reviewed used T1-weighted (T1w) pulse sequences, which give good soft tissue contrast. Sometimes, T2-weighted (T2w) sequences or a combination of T1w and T2w images are used, as T2w images offer a different type of contrast and can be particularly useful for instance to visualize and segment cerebrospinal fluid, which in turn may improve the accuracy of the reconstructed outer cortical surface for example (for an overview of MRI principles and sequences, see Westbrook et al., 2011). Similar to the discussion of field strength above, the spatial resolution of the pulse sequences have generally improved over time, and more recently started projects typically use ~1 mm isotropic voxels. Higher spatial resolution improves the accuracy of the measurements, particularly of smaller structures, and also allows for use of more fine-grained automated segmentation procedures, such as volumetric measurement of hippocampal subfields (Iglesias et al., 2015) and subdivisions of the cerebellum (Diedrichsen, 2006).

Multiple studies on adults have directly tested the reliability of MRI-derived measures of brain volume or cortical thickness across field strengths, scanner vendors, scanner upgrades, pulse sequences, the number of acquisitions (single vs. multiple averaged), parallel imaging, and scan sessions (Han et al., 2006; Heinen et al., 2016; Jovicich et al., 2013, 2009; Kruggel et al., 2010; Morey et al., 2010; Wonderlick et al., 2009). The studies generally conclude that these types of measurements are reliable. However, the results also clearly demonstrate that the effects of varying acquisition specifics are non-negligible. For example, in a recent study, 10 elderly subjects were scanned with 1.5T and 3T scanners of the same manufacturer and platform on the same day (Heinen et al., 2016). Brain volumes were relatively robustly measured for large compartments, including total grey matter and white matter (e.g., for FreeSurfer 5.3 (see section 6 below), mean absolute difference as % of mean volume: 1% and 2%, respectively). Nonetheless, effects of this magnitude clearly represent substantial sources of noise, or potentially systematic bias, in developmental studies of children or adolescents where annual change rates in most structures are in the 0-2% range (Tamnes et al., 2013). Furthermore, image acquisition differences may potentially have even larger effects for smaller brain compartments. Of particular importance for longitudinal studies, scan-rescan reliability has been shown to vary across brain regions, with relatively low reliability e.g. for the nucleus accumbens and the amygdala (Morey et al., 2010). On average, scan-rescan reliability is proportional to the volume of a structure (Morey et al., 2010) and improves when using longitudinal analysis pipelines (Jovicich et al., 2013; Reuter et al., 2012).

The general recommendation is thus that it is highly important to consider all of these image acquisition variables in both the design and analysis of longitudinal studies. Advances in this dynamic field will continue to offer opportunities to optimize these variables in order to address specific research questions. However, the implementation of novel approaches can also be problematic for longitudinal studies that must place a premium on consistency over the course of the study. One should as far as possible strive for uniformity in image acquisition within a given study, and if this is not fully possible, e.g. due to unforeseen hardware of software changes, it is critical to try to avoid systematic relationships between image acquisition variables and the variables of interest such as age. If image acquisition parameters do vary across scans, the inclusion of redundant scans that differ only in terms of these parameters can help to partially address possible confounds in a statistical model.

## 5. Quality control procedures

Another aspect of data processing that is rarely reported is the procedure used to assess image and measurement quality. Anecdotally, there appears to be much variation within the field. This is not specific to neuroimaging, as certain practices evolve and are only widely adopted after systematic testing. For example, rigorous motion control procedures in resting-state functional connectivity studies were widely adopted after the publication of several reports illustrating the impact of motion on resulting inferences, including developmental differences (Power et al., 2013, 2012; Satterthwaite et al., 2013).

Data quality can be assessed at different stages in a structural MRI study, as recently outlined by Backhausen and colleagues (2016). In addition to checking data quality at the scanner console after running a structural sequence, which can allow for re-acquisition if needed, it is critical that data quality is assessed after processing images; even acceptable raw images can fail the processing stage. For example, one study found that almost half of a large number of scans showed cortical reconstruction errors within the anterior temporal cortex (Mills et al., 2014b). Data quality can be assessed manually or by outlier detection after quantification of structure. When a scan is considered to “fail” the processing procedure, it is possible to manually intervene and reprocess the image. Certain software packages (i.e., FreeSurfer) provide extensive detail on multiple methods to do so. Nevertheless, what degree to intervene, and how to intervene, is at the discretion of the researcher, and often these details are not included in manuscripts. Assessment of the quality of processed scans remains subjective, and studies vary in their methods employed and details reported. Given the current lack of reporting of QC procedures, it is hard to fully understand their impact on resulting developmental trajectories of anatomical brain measures.

One of the most common artefacts in structural brain imaging is motion-induced artefact. Motion can be identified in a raw anatomical image by visual inspection (i.e. for ringing or waves at the periphery of the brain), and can also be systematically quantified based on predefined criteria. In addition, there are now automated Brain Images Database Structure (BIDS) apps that will assess raw anatomical MRI scans for quality and output quantitative measurements (Gorgolewski et al., 2017). However, there are no set standards for assessing motion artefact or clear cut-offs for when to consider an anatomical scan unusable. Already 15 years ago, the issue of how head motion during image acquisition could relate to anatomical brain measures in developmental neuroimaging studies was addressed in a study by the NIMH CPB (Blumenthal et al., 2002). Assessing the relationship between image quality (with motion artefact rated as “none”, “mild”, “moderate” or “severe”) and brain volume measures, findings revealed that quality was negatively correlated with age and grey matter volumes. The authors cautioned that even minimal motion artefact had a significant impact on anatomical estimates.

The same conclusion was drawn by a larger systematic investigation of head motion artefacts in MRI scans from adult participants conducted more than a decade later. Reuter et al., (2015) assessed the impact of visual inspection QC procedures on reducing motion-related artefact bias by categorizing scans as “pass”, “warn”, or “fail”. A negative relationship between motion and grey matter volume remained even after “fail” scans were removed, suggesting that QC procedures that only exclude scans with blatant motion artefact do not adequately remove the confound of motion. However, the relationship between motion and grey matter volume was no longer significant when “warn” scans were also removed, suggesting that bias from motion artefact can be more appropriately dealt with when more stringent QC procedures are implemented. This study also collected navigator images at each TR during the scan, which allowed quantification of the actual amount of head motion. Findings revealed that motion reduced estimates of grey matter volume and thickness across the majority of the cortex, with some regional variability (including increased thickness in certain regions). Even small amounts of motion introduced a spurious result of up to 2% grey matter volume loss. Further, although the relationship between motion and cortical thickness estimates was significant in images processed using FreeSurfer’s longitudinal pipeline, the relationship was even greater when images were treated as independent from one another (processed with the regular pipeline).

The impact of motion has been further confirmed and extended by recent papers using fMRI motion in the same scanning session as a proxy for motion during structural scans (Alexander-Bloch et al., 2016; Pardoe et al., 2016; Savalia et al., 2017). This proxy measure shows high inter-scan reliability and reasonable convergence with visual inspection for motion artefact in structural scans, supporting its use when explicit measurements of motion during structural scans are not available. Similar to visual inspection, scans with increased motion estimated using this proxy measure appear to exhibit decreased total brain and regional grey matter volume (Alexander-Bloch et al., 2016; Pardoe et al., 2016; Savalia et al., 2017); decreased lobar (Alexander-Bloch et al., 2016) and vertex-level thickness (Pardoe et al., 2016; Savalia et al., 2017); and increased lobar estimates of cortical curvature (Alexander-Bloch et al., 2016). There is significant regional heterogeneity in the impact of motion, which may partially result from heterogeneity in the amount of physical motion itself related to differential distance from physiological axes of rotation. Motion may impact automated morphological estimates at least partially by decreasing grey-white tissue contrast (Pardoe et al., 2016). While there does appear to be broad similarities across image processing software to the extent that this has been tested, specific differences across platforms have also been reported, such as increases in thickness in medial occipital lobe in scans with high motion artefact in FreeSurfer (Alexander-Bloch et al., 2016; Pardoe et al., 2016) but not in CIVET (Alexander-Bloch et al., 2016). This begs the question of which measurement of fMRI motion is used as a proxy, as well as exactly how scans are assessed to be qualitatively motion-free, underscoring the need for transparency and consistency in these areas.

The issue of motion-related bias is particularly problematic for developmental studies, given evidence that younger individuals on average show higher in-scanner motion than older individuals (Power et al., 2012; Satterthwaite et al., 2013). The potential impact of this issue on the longitudinal imaging literature is unclear. The effect size of minimal motion on cortical thickness measurements appears to be relatively small compared to the effect size of age in developmental populations (Alexander-Bloch et al., 2016). Based on evidence for decreasing motion with age, and decreasing cortical thickness across the second decade of life, it is unlikely that motion would account for putative reductions in cortical thickness in adolescence. However, it is unclear what the impact of motion could be on developmental trajectories starting in childhood. It can thus not be ruled out that motion-related bias could be implicated in previous reports of increases in cortical thickness through late childhood and reports of regionally and developmentally heterogeneous cortical thickness peaks. This hypothesis is supported by a recent study investigating the impact of QC procedures on developmental trajectories of cortical thickness across ages 5-22 years (Ducharme et al., 2016). While quadratic trajectories were identified when using a *standard* QC procedure (i.e. excluding scans with gross deformation of brain anatomy, large truncated brain areas, or diffuse areas of problematic grey-white boundaries definition), many of these non-linear patterns were no longer present when using a *stringent* QC procedure (i.e. excluding scans with localized areas of imprecise cortical definition, inclusion of white matter within cortex, or vice versa; see Figure 2).

**Figure 2.**
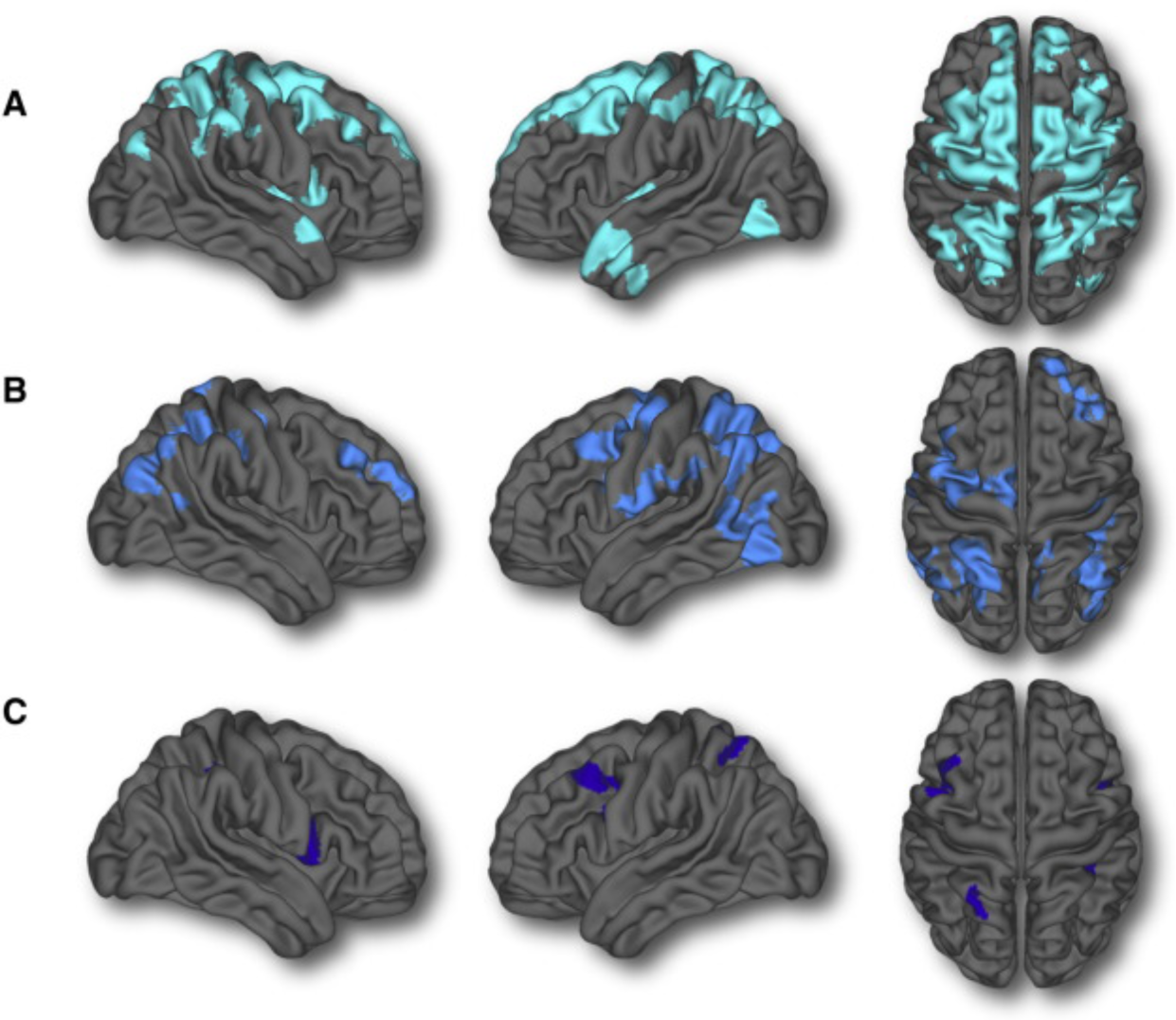
Ducharme et al.'s (2015) investigation of nonlinear developmental trajectories at different levels of quality control. Greatest quadratic or cubic trajectories (areas highlighted in different shades of blue) were evident with (a) no quality control, followed by (b) standard quality control. In comparison, minimal nonlinear trajectories were identified when (c) employing stringent quality control.

In general, scans with frank, qualitative motion artefact have greatly increased impact on morphometric estimates; but critically, motion-related bias appears to persist even within scans that are qualitatively free of motion. However, there is currently no standard, agreed-upon, protocol for QC of structural scans. Backhausen et al., (2016) recently proposed one such protocol that shows promise, recommending that the degree of post-processing QC be determined by the rating of pre-processed images. The adoption of such protocols will help increase transparency and reporting in manuscripts, which is currently lacking in the field as evident from the studies reviewed in this paper. Indeed, several reviewed studies did not include any description of QC. Others tended to focus on the quality of either raw or processed images, but rarely both (e.g., Cao et al., 2015). Limited detail was also provided about the criteria employed and manual corrections undertaken. As an exception to this, a couple of studies provided detailed information, including figures, about their QC procedures in supplementary materials (e.g., Ducharme et al., 2016; Raznahan et al., 2014). Such increased transparency and reporting in manuscripts will help us further our understanding of how different QC methods might impact the results of developmental structural imaging studies.

## 6. Image processing strategy

There are several programs available for processing anatomical brain images for morphometric analysis (see Table 2 in Mills and Tamnes, 2014). While earlier studies used a variety of manual and automated tools (Table 1), CIVET (http://www.bic.mni.mcgill.ca/ServicesSoftware/CIVET) and FreeSurfer (http://surfer.nmr.mgh.harvard.edu/) are the most commonly used contemporary automated software packages (the former used in 4/34 studies and the latter used in 16/34 studies). FreeSurfer is an openly available software package that provides global, regional and vertex-wise estimates of several brain measures. It offers a longitudinal processing stream that generates a within-subject template to increases reliability and statistical power (Reuter et al., 2012; Reuter and Fischl, 2011). However, this processing stream was developed for adult populations, and thus assumes intracranial volume (ICV) is stable in the participant across time, which is a potential drawback given that there is evidence that ICV continues to increase up to mid-adolescence (Mills et al., 2016).

CIVET provides conceptually similar anatomical estimates to FreeSurfer, but does not currently have a longitudinal pipeline. This is an important consideration given that it has been shown that using longitudinal pipelines may change developmental trajectories by reducing noise (i.e., variance) by taking advantage of the longitudinal consistency of the data (Aubert-Broche et al., 2013). Other less frequently used programs that facilitate longitudinal analysis include QUARC (Quantitative Anatomical Regional Change; Tamnes et al., 2013) and the LL Method (Aubert-Broche et al., 2013), which use a within-subject template for registration purposes alone and thus allows for variation in head size over time. Nevertheless, FreeSurfer’s (v5.3 onwards) longitudinal pipeline is the only one thus far that can be applied to scans from participants with single time points, thus ensuring consistent processing of all images used in analyses such as multilevel modelling.

As discussed above, there are acquisition methods that can be used to attempt to deal with hardware/software upgrades in longitudinal studies. While multi-site projects have used site as a covariate in analyses to account for potential biases, post-acquisition processing steps have also been applied. For example, FreeSurfer’s longitudinal stream and the LL Method have been used to assess potential scanner upgrade bias by examining change in subsets of participants before and after upgrade (Aubert-Broche et al., 2013; Dennison et al., 2013; Mutlu et al., 2013; Vijayakumar et al., 2016). Methods have ranged from calculating Cronbach's alpha or change values at the vertex level (Aubert-Broche et al., 2013; Mutlu et al., 2013), to estimating test-retest reproducibility errors for individual ROIs and testing whether the amount of change observed in the study population was likely to have occurred over and above those expected from upgrade effects alone (Dennison et al., 2013; Vijayakumar et al., 2016).

In summary, while some work has been done to assess the effects of different software on age-related differences in structural brain measures (Walhovd et al., 2016), further investigation is needed to assess such effects across the full range of software packages and versions, and also with longitudinal data. Of note, recent work has re-analyzed existing longitudinal datasets using a single software package (Mills et al., 2016). While this is a positive step in elucidating potential software differences, other factors contributing to differences between studies make it difficult to assess the influence of different software packages (and versions) on current findings.

## 7. Statistical analyses

### 7.1 Analytic methods

Given the longitudinal nature of data collected in this field, statistical analyses need to appropriately model interdependencies of observations within subjects. While there are multiple different methods to do so, most studies have employed multilevel modeling (MLM; also referred to as mixed-effects models), including 21 of the 34 studies reviewed in Table 2. MLM is particularly suited to ALD studies that collect data from individuals at different ages and differing time intervals, which in combination with missing data, result in unbalanced datasets. MLM is able to handle all available data in these instances, and consequently increases power to detect developmental effects (Gibbons et al., 2010; Singer and Willett, 2003; Verbeke and Molenberghs, 2000; West et al., 2006). Furthermore, missing observations in SCD studies means that the final dataset might still be unbalanced in nature, thus highlighting the value of this methodology (e.g., Vijayakumar et al., 2016). Newer studies are beginning to employ more flexible approaches to modelling, such as spline modeling (e.g., Alexander-Bloch et al., 2014; Tamnes et al., 2013), which might provide better fit of the underlying data by stitching together several basis functions that best fit segments of the developmental span of interest (Reiss et al., 2014; see Figure 3 for an illustration of these trajectories).

**Figure 3.**
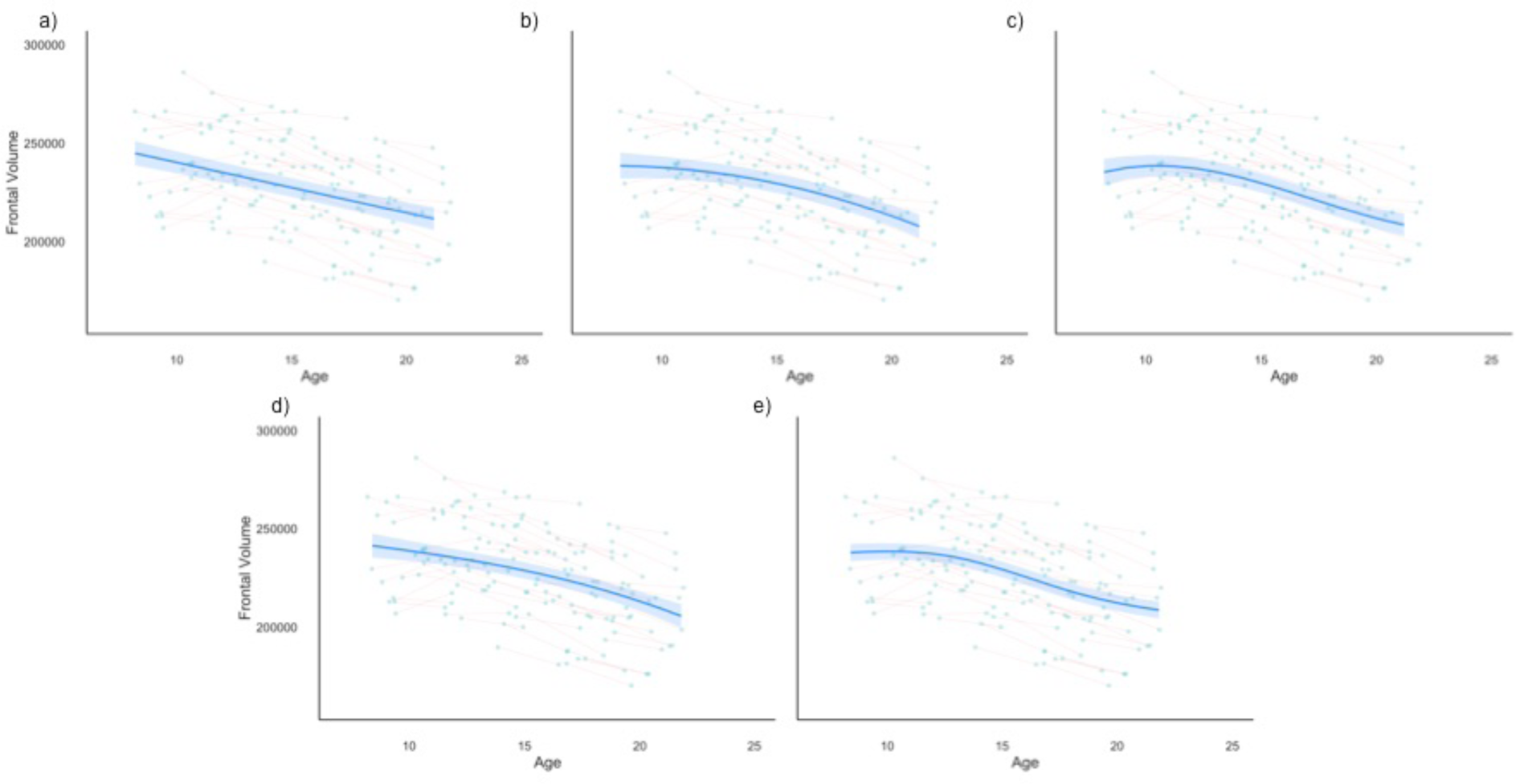
Frontal lobe volume of the Neurocognitive Development sample (Tamnes et al., 2013) modelled using different polynomial and spline modelling techniques. Figures represent *a)* linear, *b)* quadratic, and *c)* cubic polynomial trajectories, as well as spline modelling with *d)* three and *c)* five knots.

A small number of studies have chosen to calculate a change (i.e., difference or percentage change) score and conduct a single sample t-test on this index of development (5 out of 34 studies; e.g., Sowell et al., 2004; van Soelen et al., 2012). In ALD studies, this approach has also been used to examine whether age is associated with calculated change indices (Tamnes et al., 2013), thus providing valuable information about the amount of change occurring at different ages. However, the value of general linear models decreases with increasingly complex samples with multiple waves of assessments (i.e., beyond a two wave study), variation in timing of assessments, and missing data. Therefore, it is not surprising that most studies have chosen to employ MLM, which will be the predominant focus of this section.

### 7.2 Trajectories and peaks

MLM uses a likelihood-based approach to statistics that provides information about the *relative* usefulness of a model to describe data in comparison to another model. It does not, however, provide information about the absolute worth of any given model. Consequently, findings are influenced by the set of models chosen a priori to be investigated. The developmental trajectories modeled in any given study are generally based on the study design and nature of the observations. While ALD studies can examine complex non-linear trajectories at a group-level due to variance in participants’ age during assessments, modeling in SCD designs is limited by the number of repeated assessments per individual. Trajectories examined by any given study are often also influenced by prior research and prevalent theories in the field. As such, there can be a tendency for studies to examine first- and higher-order polynomial models, arising from early studies in this field identifying nonlinear patterns of brain development (Giedd et al., 1999; Gogtay et al., 2004; Shaw et al., 2008). This is evident in 15 of the 21 studies listed in Table 2 that employed MLM.

Polynomial models are popularly employed because they are able to give a rough approximation of the pattern of change in a dataset with few within-participant observations. The simplest first-order model with a linear age effect enables us to determine whether a brain measure is decreasing or increasing across development. Higher-order models impose inflection points, with a quadratic term creating a U or inverted-U pattern, and a cubic term creating an S shaped curve (see Figure 3 for an illustration of these trajectories). These inflection points have been used to provide a point estimate for when a certain brain measure “peaks.” However, there are several limitations to this procedure. “Peaks” or inflection points associated with nonlinear trajectories are often statistically reported and interpreted through solving higher-order age functions, as evident in 11 of the studies reported in Table 2.

Inflection points are theoretically appealing, being commonly interpreted as sensitive periods characterized by significant brain development. However, there is a possibility that inflection points are an artefact of the modelling strategy or age range studied (Fjell et al., 2010), as opposed to a true effect in the data. Varying “peaks” have been identified within the same dataset when studies use differing inclusionary criteria and thus report on differing subsamples from the same project. For example, The NIMH CPB sample has reported different peak ages for frontal grey matter volume, with the group reporting younger peaks as the dataset grew in sample size over the years: Giedd et al. (1999) 12.1 years in males and 11.0 years in females; Lenroot et al. (2007) 10.5 years in males and 9.5 years in females. Differences in time intervals between scans are also likely to influence identified “peaks”, as shorter time intervals between scans might be more sensitive to subtle non-linear development that might not be evident using longer intervals. Studies should be mindful of these issues when discussing peaks, and at the very least, report confidence intervals around these point estimates (e.g., Raznahan et al., 2014).

### 7.3 Model selection

In order to choose between different polynomial trajectories, two main strategies have been employed. Many of the seminal early studies used a top-down approach whereby the most complex developmental model was chosen to describe the data based solely on the significance of the polynomial parameter (Giedd et al., 1999; Gogtay et al., 2004; Lenroot et al., 2007). However, more recent studies have used model fit indices or likelihood ratio tests to ensure the most parsimonious model is selected (i.e., choosing a less complex model when the addition of parameters does not improve model fit (e.g., Mills et al., 2016; Mutlu et al., 2013; Vijayakumar et al., 2016). These different approaches may contribute to some of the variation in results, as non-linear developmental trajectories for certain measures (i.e., cortical thickness) identified in studies using the top-down approach have not consistently been replicated in more recent studies employing model-fit indices (e.g., linear trajectories identified by Ducharme et al., 2016; vs. predominantly cubic trajectories identified by Shaw et al., 2008). However, we note that given these studies differ on a number of parameters, it is not possible to specifically attribute variation in results to the issue of model selection alone.

Traditionally, MLM does not give preference to a specific model selection approach. Nevertheless, top-down approaches are best used when there is a strong theory to guide them (see King et al., this issue). But a problem can arise when guiding theories, themselves, are based solely on top-down approaches that bias results to more complex models. This is evident in our field, where the predominant theories about anatomical brain development (e.g., nonlinear trajectories) were inferred from studies using a top-down approach. This had a subsequent ripple-down effect as latter studies also employed similar top-down approaches, thus perpetuating the issue. In total, we identified 12 studies that used the top-down approach for selecting between different age-related trajectories, in comparison to only 5 studies using model fit indices. Nevertheless, there does appear to be a shift with more recent studies increasingly employing model fit indices into their investigations on brain development, and we need to strive to incorporate these findings into our theories in order to progress the field. One way to compare across studies using different model selection criteria would be to reduce emphasis on the actual model fit (i.e., cubic, quadratic or linear) and instead focus on the overall pattern of change. This would, for example, emphasize similar periods of stability/change between quadratic and cubic trajectories, or similar overall direction of change between first- and higher-order polynomial trajectories. Further, the inclusion of confidence intervals lessens the stark differences between different polynomial trajectory shapes, and highlights the lack of specificity that can be derived from model fits (e.g., Mills et al., 2016).

### 7.4 Group vs individual differences

Interestingly, all the identified studies in Table 2 that employed MLM only obtained group-level (i.e., fixed effect) developmental trajectories. Although subject-level trajectories can be modeled as random effects to account for interdependencies in the data (Pinheiro and Bates, 2013; Verbeke and Molenberghs, 2000), only one study was identified that employed this technique (Aubert-Broche et al., 2013), with all others only incorporating a random effect for the subject (i.e. random intercept). Information regarding differences in model fit indices when incorporating random slopes would provide valuable information about variability between individuals in their trajectories. Furthermore, studies considering cross-level interactions between random and fixed effects would provide novel insight into whether individual differences interact with group-level heterogeneity in meaningful ways (e.g., Ordaz et al., 2013). Most studies have examined the interaction between age and sex in predicting brain structure, for example, but the fixed effect of age has exclusively been used in this interaction term, thus only utilizing information about group-level differences in the *age* term. Such group-level analysis is often necessitated in ALD designs that are interested in modeling complex developmental trajectories that cannot be examined at an individual level (i.e., cubic trajectories modeled despite some individuals only having three or less repeat assessments).

A small number of studies have tried to address this issue of intra-individual variability by describing both group- and individual-level change in their sample. For example, 3 out of the 34 identified studies in Table 2 graphically illustrate the percentage of individuals that exhibit increases, decreases, or no change, in a structure over time. Dennison and colleagues (2013) apply this technique to a SCD study, whereas Lebel and Beaulieu (2011) and Zhou et al. (2015) bin their subjects into age groups given the ALD nature of their sample. It is also beneficial to report the magnitude of change, which has been addressed in some studies using plots of raw within-subject change over time. However, the usefulness of this strategy can also vary based on other study characteristics (e.g., difficult with vertex-wise as opposed to region of interest analyses).

Analyses of difference scores (e.g., annualized percentage change), described above, are also generally conducted at the group level. However, reporting variance in these measures, along with means, would provide some indication of the level of individual differences in a sample. To our knowledge, none of the current studies have done so. Percentage change scores are also valuable when interpreting the association between different developmental variables (i.e., how are individual differences in development on a particular measure associated with change in another measure). For example, one of the reviewed studies employed difference scores to qualitatively explore, through graphical illustration, the relative contributions of changes in thickness and surface area to volumetric development (Raznahan et al., 2011b).

In summary, most studies fail to address the issue of individual differences in brain development, and those that do so use different strategies, each with their own benefits and limitations. While it is not possible to recommend one particular strategy, it is important that studies provide some report of individual variability in their sample. The role of individual differences will also become increasingly important as attention turns towards understanding interactions between group-level characteristics (e.g., environment, genes) and intraindividual variation in trajectories.

### 7.5 Specificity of analyses

Statistical analyses can be conducted at differing levels of spatial specificity, varying from global to voxel/vertex-level indices. Between these two extremes lies the lobar- and parcellation-based approaches, which groups voxels/vertices into regions based e.g. on anatomical landmarks. A review of studies in Table 2 revealed roughly similar distribution of these approaches, with 10 studies each using global, lobar and vertex-wise methods and 11 studies using parcellations. Note that many studies chose to employ more than one approach, in addition to a smaller subset that employed region-of-interest analyses. While voxel/vertex-level analyses provide the greatest spatial resolution, the parcellation-based approach can be easier to interpret given that uniform developmental patterns are attributed to each structurally homologous region, thus providing a middle ground between vertex-level and lobar/global measures. Vertex-level analyses typically employ a smoothing kernel to reduce noise, and prior research has shown that the size of kernels can impact on scan-rescan estimates (Han et al., 2006). However, the majority of studies do not report on the size of this kernel. While parcellation-based approaches do not require these smoothing procedures as boundaries are already defined based on anatomical landmarks, there are several parcellations to choose from, also with differing spatial resolutions (and associated boundary-defining procedures; e.g., FreeSurfer’s Desikan-Killiany vs Destrieux atlases vs Human Connectome Project’s multimodal parcellation).

It is standard that vertex-level analyses involve correction for multiple comparisons, given that analyses are conducted across tens of thousands of data points, using procedures such as false discovery rate, random field theory or Monte Carlo simulations. However, these tend to be conducted on p-values of a single model, as opposed to model fit indices. On the other hand, studies using parcellation-based data have tended to focus on model-selection procedures without correction for multiple-comparisons (as evidenced by all but four of the studies in Table 2 that employed this approach), even though the coarsest parcellation map typically outputs a large number of different regions to test. While this approach might be acceptable given that likelihood based analyses are theoretically distinct from null-hypothesis significance testing, at the very least, these findings need to be discussed carefully so that they are interpreted in terms of relative evidence as opposed to absolute conclusions (i.e., without reference to the comparison models). Plotting of effect sizes across different parcellations would also provide complementary information about the degree of change across the brain. Multivariate analyses that include all parcels within the same model are promising approaches that overcome the issue of multiple comparisons (Ziegler et al., 2016).

### 7.6 Statistical programs

There are a number of software programs available to conduct statistical analyses – either specialized or not – for neuroimaging data, and each with their own strengths and weaknesses. FreeSurfer provides easily accessible code for general linear modeling, and also computes change metrics (i.e., annualized percentage change) that can be used within such a framework. While FreeSurfer does not incorporate MLM into its main platform, there is a separate Matlab toolbox to support these analyses. SurfStat is another Matlab toolbox supporting MLM that was originally created for use with CIVET-processed data. Both toolboxes provide easily accessible methods to conduct MLM at a vertex-wise level across the cortical surface, including tools to correct for multiple comparisons (i.e., false discovery rate and random field theory correction). However, neither program supports the use of model fit indices to ascertain the best-fitting developmental trajectory. Rather, these programs are limited to a top-down approach for model selection.

In order to overcome this limitation, one of the reviewed studies employed both vertex-level top-down model selection and parcellation-based indices of fit selection, finding similar developmental maps with both approaches (Vijayakumar et al., 2016). Other studies have employed MLM within general statistical packages (i.e., *nlmefit* in Matlab and *lme4* or *nlme* in R) to conduct vertex-level analyses with a model-fit approach, with one study additionally correcting for multiple comparisons across the cortical mantle following model selection (i.e., nlmefit in Matlab followed by Monte Carlo simulations in FreeSurfer, Mutlu et al., 2013). Thus while it is possible to overcome these software limitations using a combination of different approaches and programs, the development of statistical programs specific to longitudinal neuroimaging analyses with more sophisticated options would be a welcome addition to the field (see Madhyastha et al., this issues).

## 8. Treatment of important covariates

### 8.1 Sex differences

The majority of studies in this field have investigated sex differences in developmental trajectories of brain structure (evident in 22 out of the 34 studies in Table 2), while a smaller number (3 out of 34 studies) have chosen to instead control for sex in their analyses. Almost all studies investigating sex differences have identified developmental trajectories across the sample prior to investigating whether the incorporation of sex main effects or interactions with age improve model fit. However, this approach precludes the identification of different trajectories (e.g., linear versus quadratic) between the sexes, and assumes that males and females exhibit similar developmental patterns (e.g., both present with linear growth, although rate of growth can differ). Furthermore, if differing developmental patterns do exist, identification of one developmental trajectory for the entire sample might not be truly reflective of either sex. In order to overcome this issue, some studies have chosen to analyze males and females separately (e.g., Goddings et al., 2014). However, these are limited to discussing qualitative differences between the sexes. Future studies may benefit from combining these approaches, as implemented by Lebel and Beaulieu (2011), such that each sex is first examined separately and only combined if similar trajectories are identified (to examine development of the group as a whole, as well as potential sex differences).

Research on sexual dimorphism in the brain is also influenced by if, and how, studies account for differences in global brain size. Males exhibit larger global brain sizes than females from childhood through to adulthood (Giedd et al., 2012; Jahanshad and Thompson, 2017; Paus et al., 2017). Therefore, studies attempt to control for overall brain size to reduce the risk of observing structural brain differences between sexes that is solely due to differences in overall brain size. These methodologies, and their impact on sex differences, are further discussed in the following section.

### 8.2 Whole brain correction

As highlighted in Table 2, some studies choose to account for global brain size in their analyses, but many choose to not do so. There are a number of issues faced by researchers when deciding to correct for global brain size, including potential measures to use and methods to employ. When considering methodology, most studies have included global brain size as a covariate in statistical analyses (6 out of 34 studies reviewed), though some have chosen to correct using the proportion method that divides regional size by global size (5 out of 34 studies reviewed). The former covariate method is often preferred for vertex-wise analyses as it is easier to implement in comparison to the latter approach, which would require calculation of adjusted brain measures across the cortical mantle prior to statistical analyses. However, an important limitation of both these popular methods is the assumption of linear scaling between regional and global brain size.

Scaling factors in the brain are constrained by metabolic and physical principles, such that neuronal size and other components of brain anatomy undergo a non-uniform enlargement of subcomponents with increasing overall size (Toro et al., 2009). These nonlinear relationships between structure and size, referred to as allometric principles, are demonstrated by exponential increases in white-to-grey matter ratios at a rate of 4:3 with increasing total brain volume (Zhang and Sejnowski, 2000). This might account for greater grey-to-white matter ratio in females due to their smaller overall brain size, supported by findings of minimal sex differences in this ratio when accounting for differences in overall brain size (Leonard et al., 2008). On a more regional scale, cross-species work has shown nonlinear scaling of subcortical volume with increasing whole brain volume (WBV; Finlay and Darlington, 1995), which was confirmed by a recent investigation in humans (Reardon et al., 2016). Furthermore, violations of these allometric principles by commonly used proportion and covariate correction methods confounds the effects of nonlinear scaling and group effects (i.e., sex) on subcortical volume (Reardon et al., 2016). Greater consideration of regional allometric scaling laws may therefore provide greater clarity into the specificity of findings on sex (and other group) differences.

Another important issue for developmental neuroimaging is that global brain size continues to change during adolescence, and differences in development rates across the brain could bias results when controlling for global size. Moreover, group differences in global measures could be a reflection of earlier maturation in one group, thus potentially eliminating differences of interest when it is controlled. In order to deal with this problem, some studies have controlled for ICV as initial research suggested it stabilized between early and mid-adolescence (Courchesne et al., 2000; Pfefferbaum et al., 1994). Specifically, three out of 10 studies that chose to control for global brain size in Table 2 used ICV. In comparison, 8 of these studies controlled for WBV (including one that employed both measures). However, as mentioned above, Mills and colleagues (2016) found that both ICV and WBV continued to develop during adolescence. Furthermore, controlling for these two measures influenced the resultant regional (i.e., grey vs white matter volume) trajectories differentially, and these two measures had varying impacts on sex differences based on the correction method employed. While both ICV and WBV accounted for sex differences when using the proportion method (i.e., the addition of sex to proportion-corrected models did not improve model fit), WBV alone was able to do so with the covariance method. Considered within the context of current findings in the literature, these results suggest that the estimate of global brain size employed by studies likely influenced the sex effects that were observed. Therefore, it is considered best practice to present both raw and corrected brain measures when examining the influence of sex on brain development (e.g., Dennison et al., 2013). Furthermore, to our knowledge, there has been no investigation into the effects of controlling for change in global brain size. While the scaling factors discussed above will still impact this methodology, it might be a valuable area for investigation as it accounts for continued changes in global brain size over development.

Global volumetric estimates are sometimes also used to control for whole brain size in analyses of non-volumetric measures (i.e., cortical thickness). This approach is not without fault as volume is driven by both thickness and surface area, with some research suggesting that it is largely driven by surface area (Im et al., 2008; Raznahan et al., 2011b). Only minor change has been identified in cortical thickness with enlargements of brain size, consistent with theoretical models by Van Essen (1997) and Rakic (1988). These findings and theories question the influence of increasing brain size on cortical thickness. On the other hand, matching the global measure with the metric of interest, particularly in relation to cortical thickness (i.e., controlling for average cortical thickness), has also been questioned given wide variability in thickness across the cortex (Palaniyappan, 2010). Therefore, the majority of research on non-volumetric measures have chosen not to control for whole brain size.

Given these issues, due consideration needs to be given when choosing the method of correction/control and measure of global brain size, as well as interpretation of findings. Most researchers in this field are aware of these issues, and thus present uncorrected results if they choose to control for global brain size. However, as highlighted by Mills and Tamnes (2014), it would also be valuable to present the developmental effects of the global measure used, so that readers can fully understand whether results were driven by global or regional changes. Furthermore, investigation into developmental scaling relationships between global and regional measures of interest, as well as potential group differences, will help us better understand the influence of correction procedures and allow us to more appropriately interpret findings.

### 8.3 Pubertal maturation

While most studies on brain development have indexed maturation using age, a growing number of studies are examining the influence of puberty. This interest partly stemmed from early studies identifying earlier “peaks” in cortical volume in females compared to males, which roughly corresponded to timing differences in the onset of puberty between the two sexes (Giedd et al., 1999; Lenroot et al., 2007). Although the exact age of these “peaks” has been found to vary significantly since these initial findings, along with many studies failing to identify any sex differences for certain measures of cortical development (e.g., cortical thickness - Mills et al., 2014b; Vijayakumar et al., 2016; Wierenga et al., 2014b), there remains a growing interest in pubertal influences. There are a number of cross-sectional studies on puberty-related cortical development, but only a handful of longitudinal studies thus far (outlined in Table 3). These studies inevitably have to consider potential age effects, as pubertal maturation and age are highly covaried (Braams et al., 2015). Three of the studies either controlled for age-effects or examined interactions between age and pubertal measures. In comparison, two studies, from the University of Pittsburgh cohort, recruited participants within a minimal age span given the project’s primary focus was related to pubertal effects on brain development. However, as there remained some variance in age in this sample, the authors chose to control for baseline age within their analytic models (Herting et al., 2014). Therefore, studies choose the most appropriate methodology to deal with age depending on their study design. However, it is always important to check for multicollinearity when entering age and puberty into the same model, as well as recognizing that controlling for age might absorb much of the variance related to puberty given the strength of correlation between these two variables. These studies might therefore benefit from presenting results both with and without the inclusion of age in their models.

Puberty-related studies also have to consider which index of pubertal maturation to investigate. Although a detailed discussion of this topic is beyond the scope of this paper (for a review, refer to Shirtcliff et al., 2009), it is interesting to note that all studies listed in Table 3 have investigated pubertal/Tanner stage using self- or parent-report measures given ease of administration and minimal costs of processing data. However, most of them have also investigated associations between cortical development and gonadal sex hormones (i.e., testosterone and estradiol), and only one study thus far has investigated how cortical development may be related to interactions between different hormones (Nguyen et al., 2013). Pubertal studies also have to deal with issues related to modeling sex differences described above, as well as sex differences in pubertal maturation. Given that females tend to exhibit physical signs of puberty 1-2 years earlier than males (Marshall and Tanner, 1969; Sun et al., 2002), projects specifically interested in pubertal development sometimes recruit younger females at baseline compared to males, with the aim of capturing pre-pubertal stages in both sexes (e.g., University of Pittsburgh project; Herting et al., 2014).

**Table 3.** Details of longitudinal studies investigating pubertal maturation in relation to structural brain development.

| Studies | N (M) | N Scans, n per subject, interval, range | Age (y) | Image processing software (version) | Pubertal measures | Measures: vol/sa/ct/others | Specificity of analyses | Index of analyses: absolute or change values, whole brain correction | Statistical analyses: analysis method (software) variables model fit trajectories multiple comparison |
| --- | --- | --- | --- | --- | --- | --- | --- | --- | --- |
| Nguyen et al., 2013a (NIH MRI) | 281 (117) | 479, 1-3 per subject, 2 year interval | 4 - 22 | CIVET | Stage Testosterone | CT | Vertex-wise | Absolute | Mixed models: SurfStat<br>Effects: puberty (testosterone, stage), age, sex and hemisphere interactions, controlling for WBV, testosterone collection interval<br>Trajectories: linear<br>MC: RFT |
| Nguyen et al., 2013b (NIH MRI) | 255 (112) | 407, 1-3 per subject, 2 year interval | 4 - 22 | CIVET | Stage Testosterone DHEA | CT | Vertex-wise | Absolute | Mixed models: SurfStat<br>Effects: 1. DHEA, age and sex interactions, controlling for WBV, salivary collection times, scanner, handedness; 2. DHEA, testosterone and sex interactions.<br>Trajectories: linear<br>MC: RFT |
| Goddings et al., 2014 (NIMH CPB) | 275 (158) | 711, $\geq 2$ per subject, 2 year interval | 7 - 20 | FreeSurfer 5.1 | Stage | Volume | Segmentation | Absolute and percentage change | Mixed models: R<br>Effects: pubertal stage, age, and interactions (separate models for males and females)<br>Model selection: step-down for age, LRT/AIC for age*puberty models<br>Trajectories: linear, quadratic, cubic |
| Herting et al., 2014 | 126 (63) | 162, 2 per | 10 - 14 | FreeSurfer 5.1 | Stage Testosterone | CV | Global ROIs | Absolute | Mixed models: R<br>Effects: age, sex, puberty, and interactions, |
| UPitt |  | subject,<br>2 year<br>interval |  |  | Estradiol |  |  |  | controlling for ICV<br>Model selection: LRT and AIC<br>Trajectories: linear |
| Herting et al.,<br>2015<br>UPitt | 81 (33) | 162,<br>2 per<br>subject,<br>2 year<br>interval | 10 - 14 | FreeSurfer 5.1 | Stage<br>Testosterone<br>Estradiol | CT, SA | Vertex-wise | Average and<br>percentage change | Linear regressions: FreeSurfer<br>Effects: pubertal change, sex, and<br>interactions, control for baseline age,<br>puberty (and scan interval for models<br>predicting average measures)<br>Trajectories: linear<br>MC: Monte Carlo simulations |

NB: Studies are grouped by project, and subsequently ordered by year published and author surname. AIC = Akaike Information Criteria ; CT = cortical thickness ; CV = cortical volume; DHEA = Dehydroepiandrosterone; ICV = intracranial volume; LRT = likelihood ratio test; NIH = National Institute of Health; NIMH CPB = National Institute of Mental Health Child Psychiatry Branch; ROI = region of interest; RFT = Random Field Theory; SA = surface area; UPitt = University of Pittsburgh

## 9. Discussion

In summary, given that studies to date differ on a number of methodological issues, it is not surprising that there are variations in the normative brain developmental trajectories that have been identified. While it is beyond the scope of this paper (and perhaps impossible) to provide specific reasons as to why there is variation in reported results, we would like to point out that the evolution of methodological techniques used within and across projects over time is beneficial to the field. Although not to belittle the methods and results from earlier studies, the more recent papers have implemented certain strategies that are now widely accepted as improvements in our practices. For example, there appears to be a shift towards the identification of the most parsimonious models to describe the underlying data, as demonstrated by more recent studies favouring the use of model-fit indices or likelihood ratio tests in MLM. Nevertheless, there are also additional practices that would be beneficial for our field to adopt, which are outlined as recommendations in Table 4. For example, it has become apparent that many studies run independent multilevel models in regions across the brain, as defined by parcellation maps, without correction for multiple comparisons. Further consideration of this issue is required in the field, and at the very least, we need to appropriately interpret the findings of likelihood-based analyses (i.e., relative to comparison models) as opposed to a focus on p-values of each model individually.

**Table 4.**
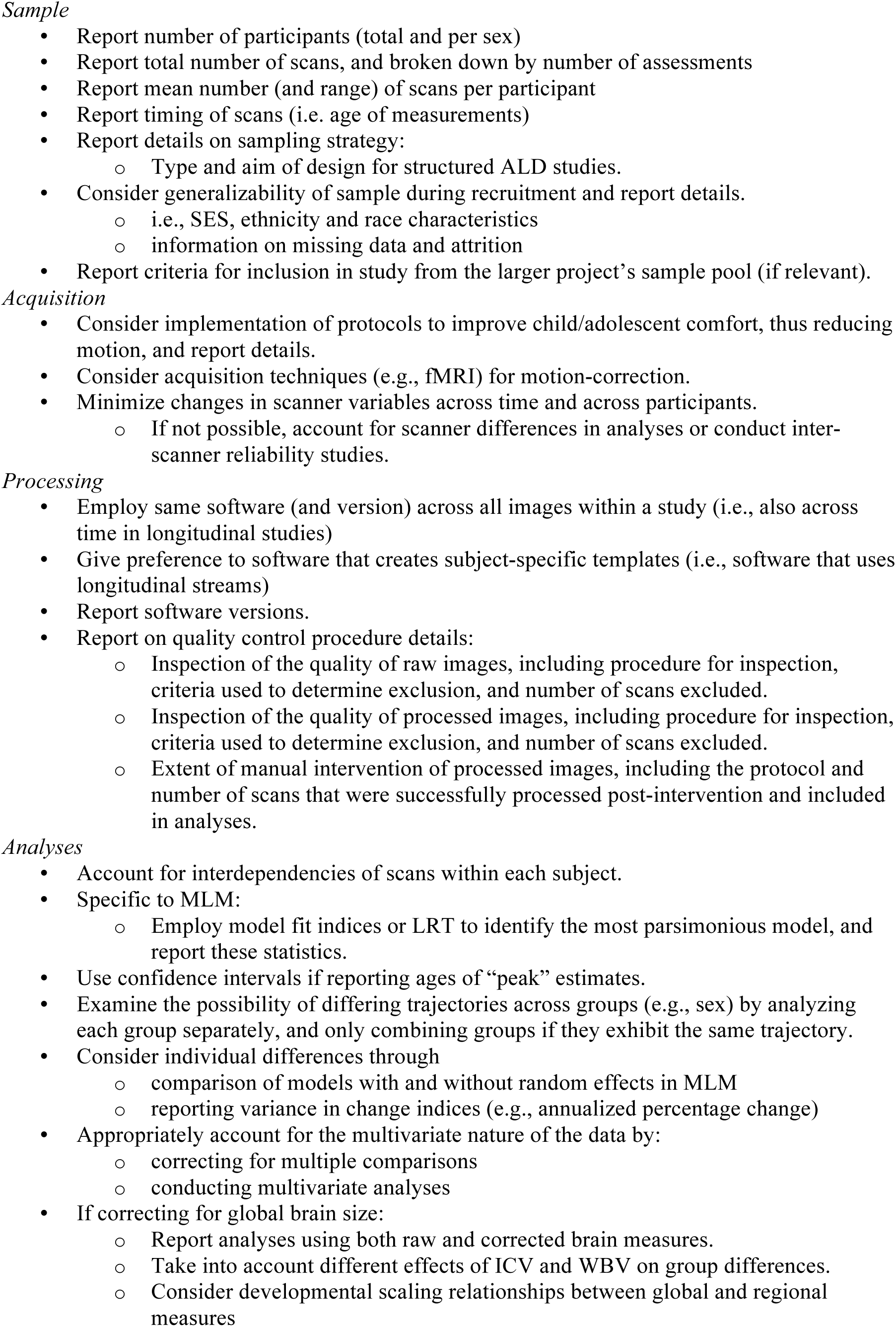
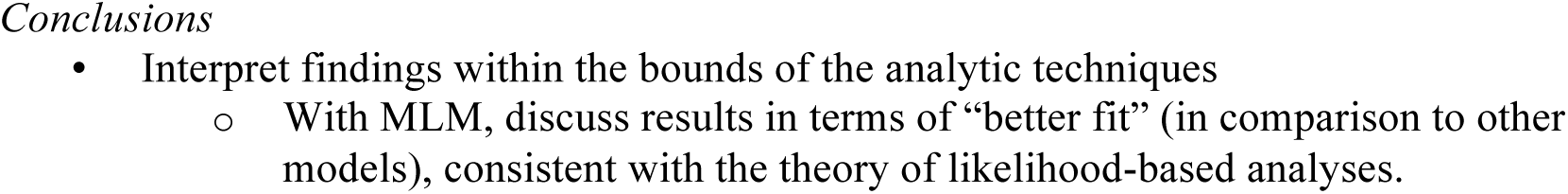
Guidelines for reporting methodological detail in longitudinal structural brain imaging studies.

This review of the literature identified certain methodological considerations that have been empirically examined, and thus it is possible to draw stronger conclusions and make recommendations regarding them. For example, there is general consensus on the benefits of using longitudinal processing streams that create subject-specific templates when reconstructing the cortical surface. Similarly, it is widely acknowledged that efforts should be made to maintain the same image acquisition parameters across subjects and time, with appropriate testing or accounting for any differences within the sample. In comparison, there has been no investigation of whether model selection procedures (i.e. step-down vs model fit indices) impact the resultant trajectories. Furthermore, although the importance of certain methodological issues has risen in prominence, there still remain no standardized practices to address many of these problems. For example, despite an increased awareness of the effect of motion and importance of QC, there are no systematic procedures that are agreed upon within the field. As such, our recommendations outlined in Table 4 are aimed towards *i)* ensuring that empirically supported ‘best-practices’ are incorporated, and *ii)* increasing transparency of other practices to support future empirical investigations and guidelines.

With an inevitable shift towards consensus regarding methodological approaches, the promise of more reliable data on normative brain development becomes likely, particularly with the ability to conduct meta-analyses. However, we are currently limited in our ability to conduct such analyses due to methodological variation. For example, it is difficult to amalgamate studies that have used parcellation versus vertex-wise approaches or raw versus percentage change analyses. One way to overcome this issue in the short term at least is for future studies to conduct replications across samples, whereby the same processing and analytic techniques are employed (e.g., Mills et al., 2016; Tamnes et al., 2017). Greater transparency and consensus regarding methods will also facilitate the implementation and interpretation of research beyond normative developmental trajectories, such as that examining the clinical and behavioral significance of brain development.

A limitation of this review is that we only included and reviewed studies assessing the development of independent grey matter structural properties (Table 2). A growing area of research in grey matter development is structural covariance, which refers to correlations across people in the morphological properties of pairs of brain regions (or larger brain networks; Alexander-Bloch et al., 2013a; Evans, 2013). Interest in this methodology lies largely in its potential to shed light on brain connectivity and connectomics using T1w scans. Indeed, there is evidence that patterns of structural covariance partially (though not entirely) recapitulate known functional boundaries (Chen et al., 2011; Li et al., 2013), resting state fMRI connectivity patterns (Kelly et al., 2012; Seeley et al., 2009) and white matter connectivity derived from diffusion MRI (Gong et al., 2012). Studies of children and adolescents have begun to map out developmental changes in structural covariance. A brief discussion of existing cross-sectional and longitudinal studies, and methodological issues specific to this methodology, is provided in Box 2.

#### Box 2

Structural covariance is an increasingly used methodology, referring to correlations across people in the morphological properties of pairs of brain regions (or larger brain networks; Alexander-Bloch et al., 2013a; Evans, 2013). While cortical thickness and grey matter volume have been the most commonly examined morphological substrates, there is evidence that other phenotypes such as cortical surface area may have specific patterns of covariance (Sanabria-Diaz et al., 2010). Studies of children and adolescents have begun to map developmental changes in structural covariance. Generally, structural covariance appears to become more widely anatomically distributed over time, but different sub-networks also follow different developmental patterns (Khundrakpam et al., 2013; Zielinski et al., 2010). For example, a study of 5-18 year-olds found grey matter covariance of primary sensory and motor areas to peak in early adolescence, while covariance patterns of regions subserving higher cognitive functions expanded linearly throughout the age range (Zielinski et al., 2010). The balance between integrative and segregative graph-theoretic properties, derived from large-scale covariance of cortical thickness between regions, was also reported to follow a nonlinear trajectory in 5-18 year olds (Khundrakpam et al., 2013). Developmental studies of structural covariance may reflect developmental changes in brain function, for example, a negative association was found between amygdala volume and prefrontal cortical thickness in children and adolescents, mirroring reports of amygdala functional connectivity (Albaugh et al., 2013).

Complementary to these cross-sectional studies, longitudinal studies have used the approach of ‘maturational covariance’ - covariance in longitudinal changes across subjects. Similar to cross-sectional studies, these longitudinal analyses have found regionally heterogeneous maturational covariance, with stronger correlations reported with association areas compared to primary sensory areas of cortex (Raznahan et al., 2011a). These statistical relationships also appear to recapitulate known functional relationships, for example, longitudinal changes in hippocampal volume were found to covary with longitudinal changes in cortical areas involved in episodic memory (Walhovd et al., 2015). Maturational covariance, reflecting coordinated development between brain regions, may in fact cause cross-sectional structural covariance (Alexander-Bloch et al., 2013b), as a generalizable biological link may hold between phenotypic covariance and coordinated maturation (Riska, 1986). In the brain, covariance patterns are likely to be established very early in development, and a recent study of children under two years old found maturational covariance to predate structural covariance (Geng et al., 2016). Notably, longitudinal studies to date have investigated the covariance of longitudinal phenotypes, as opposed to longitudinal changes in structural covariance *per se*, as covariance patterns have only been defined at the group level. Novel methodological approaches may be required to optimally leverage longitudinal data in future analyses.

In general, developmental studies of structural covariance confront an expanded array of methodological issues in addition to those of developmental anatomical imaging in general. An analogy can be drawn between functional connectivity (derived from correlations across time in brain activity) and structural covariance (derived from correlations across people in brain morphology). Comparable methodological principals should thus be applied when using multivariate analyses such as seed-based regression, principal components or graph-theoretical analyses. Brain parcellation may particularly impact covariance studies as the size of a brain region may influence its covariance patterns; we therefore advocate the use of uniformly-sized brain regions when possible. Some estimate of global effect such as total brain volume is generally also included as a covariate in studies of structural covariance, as are gender and age within experimental groups. Ideally, pending a consensus regarding appropriate statistical approaches, analyses will be presented both with and without covariates likely to impact the outcome of interest. Although specific studies of the effect of motion artefact on structural covariance have not been performed, it is likely that regions disproportionately affected by motion will manifest artefactually elevated covariance. For future research, the complexity of these methodological issues will necessitate a proportional investment in both rigor and transparency.

Moving beyond the characterization of normative brain development, many researchers are now turning their attention towards how individual differences in brain development might relate to various aspects of functioning, including cognition, affect, and behavior. While several studies have found that greater cortical thinning is related to better functioning (e.g., Ducharme et al., 2012, 2014; Shaw et al., 2006, 2011; Vijayakumar et al., 2014a, 2016), inconsistencies exist both within and across studies. While this research enables us to better understand how various developmental processes might relate to one-another, there are a number of additional methodological considerations that might be influencing these findings. A brief discussion of some of these issues is presented in Box 3.

#### Box 3

Aside from the methodological factors affecting studies of normative brain development, there are a number of additional challenges faced by studies examining how individual differences in developmental trajectories relate to cognitive, affective and behavioral development. While some studies have found that greater cortical thinning is related to better affective and cognitive functioning (e.g., Ducharme et al., 2012, 2014, Shaw et al., 2006, 2011; Vijayakumar et al., 2014a), others have found that less thinning is related to better functioning (Friedel et al., 2015). Apart from the influence of differences in the behavioral (i.e. functioning) measures employed, results of these studies are influenced by the age range of participants, as faster rates of thinning might be adaptive in certain ages but not others. Adaptive patterns of cortical development may also vary across the brain, and might be moderated by sex for certain behaviors that differentially develop in males and females (e.g., emotion regulation; Vijayakumar et al., 2014b). Development prior to the examined period might also influence identified brain-behavior associations in unknown ways. Given these caveats, care needs to be placed on the conclusions drawn from such research. It also highlights the need for further studies that replicate findings in order to confirm inferences drawn from this line of research.

From a statistical perspective, the majority of this literature has only examined behavior at one or two time points, which can be incorporated as an absolute or change (i.e., difference or residualized change) value. However, analyses will become more complex as studies incorporate three or more time points of behavioral data along with a similar number of imaging data, which would enable investigation of nonlinear patterns of correlated brain-behavior development. While MLM can handle time varying covariates along with a time varying dependent variable, it does not model change in the covariate and thus cannot examine the association between changes in two variables. MLM can examine whether the association between brain and behavior varies with age, and while this question is of no doubt of interest to many researchers, patterns of correlated brain-behavior change are also likely to provide valuable information about developmental processes. Parallel process models within the structural equation modeling framework are ideal for investigating the association between development of two or more variables, achieved via estimation of an underlying latent “change” factor for each variable and associated correlation between these factors. However, structural equation modeling is not currently supported by statistical programs for neuroimaging data, and can thus only be carried out using a region-of-interest or parcellation-based approach. Moreover, many non-imaging statistical packages do not support unbalanced datasets for structural equation modelling, and thus cannot be easily utilized on many samples without excluding a substantial amount of data. Further consideration of these factors an potential development of software packages to support this type of analyses is required for our field to progress in this area of research.

Importantly, given the observational nature of this line of research, it is not possible to comment on causality when examining brain-behavior associations; i.e., does engagement in behavior affect brain maturation or vice versa. Studies on training and practice effects provide some support for a causative role of experience and activity-dependent plasticity (Dehaene et al., 2010; Draganski et al., 2004; Gaser and Schlaug, 2003). However, it remains plausible that trophic effects influence neurobiological development that supports adaptive functioning (Burgoyne et al., 1993). Therefore, it is important that conclusions are not drawn about causality, which should rely on future animal research that attempts to fully unpack and better understand these associations.

Building on knowledge gained from research on normative brain development, a number of projects are focused on characterizing aberrant trajectories of brain development associated with psychiatric and developmental disorders or subclinical symptoms, with the aim of identifying underlying neurobiological mechanisms that might be targeted in future interventions. The majority of research has thus far concentrated on attention deficit hyperactivity disorder (ADHD), autism spectrum disorders (ASD) and schizophrenia. Studies on childhood-onset schizophrenia have identified diffuse cortical thickness differences during childhood in comparison to healthy controls, possibly with reductions in thickness becoming localized to the frontal and temporal lobes during adolescence (Greenstein et al, 2006). ASD research has produced conflicting findings, with some identifying reductions (Hardan et al., 2006), and others finding exaggerations (Wallace et al., 2010), in cortical thinning over time. Research on children with ADHD or attention problems suggest that these children have thinner cortices during childhood, and delayed or slowed thinning during adolescence (Shaw et al., 2007; Ducharme et al., 2012). The significance of these findings is limited by the same issues as have been discussed in this review. However, they are additionally limited by heterogeneity within each disorder, as well as the lack of specificity of findings to one particular disorder. Future studies on these and other disorders thought to have neurodevelopmental origins (e.g. depression, substance-use, conduct disorder) should attempt to identify whether there are consistent patterns of aberrant trajectories of brain development, and whether patterns can be differentiated across different disorders.

In conclusion, we have identified a number of methodological factors and issues, from image acquisition to data modelling, where variation in approaches taken in the current literature are likely to have contributed to differing results, and hence differing interpretations about grey matter structural brain development during childhood and adolescence. As such, it is important that results are interpreted within the context of these (and other) methodological choices. There is also wide variability in the extent to which different methodological considerations have been empirically examined. Future research should, in addition to adopting greater transparency of practices, seek to empirically examine the effects of varying methods on results, in order to promote best-practice guidelines, and ultimately, a solid and accurate understanding of child and adolescent brain development.

## References

Albaugh, M.D., Ducharme, S., Collins, D.L., Botteron, K.N., Althoff, R.R., Evans, A.C., Karama, S., Hudziak, J.J., Brain Development Cooperative Group, 2013. Evidence for a cerebral cortical thickness network anti-correlated with amygdalar volume in healthy youths: implications for the neural substrates of emotion regulation. NeuroImage 71, 42–49. doi:10.1016/j.neuroimage.2012.12.071

Alemán-Gómez, Y., Janssen, J., Schnack, H., Balaban, E., Pina-Camacho, L., AlfaroAlmagro, F., Castro-Fornieles, J., Otero, S., Baeza, I., Moreno, D., Bargall, N., Parellada, M., Arango, C., Desco, M., 2013. The Human Cerebral Cortex Flattens during Adolescence. J. Neurosci. 33, 15004–15010. doi:10.1523/JNEUROSCI.1459- 13.2013

Alexander-Bloch, A., Clasen, L., Stockman, M., Ronan, L., Lalonde, F., Giedd, J., Raznahan, A., 2016. Subtle in-scanner motion biases automated measurement of brain anatomy from in vivo MRI. Hum. Brain Mapp. 37, 2385–2397. doi: 10.1002/hbm.23180

Alexander-Bloch, A., Giedd, J.N., Bullmore, E., 2013a. Imaging structural co-variance between human brain regions. Nat. Rev. Neurosci. 14, 322–336. doi:10.1038/nrn3465

Alexander-Bloch, A., Raznahan, A., Bullmore, E., Giedd, J., 2013b. The Convergence of Maturational Change and Structural Covariance in Human Cortical Networks. J. Neurosci. 33, 2889–2899. doi:10.1523/JNEUROSCI.3554-12.2013

Alexander-Bloch, A.F., Reiss, P.T., Rapoport, J., McAdams, H., Giedd, J.N., Bullmore, E.T., Gogtay, N., 2014. Abnormal cortical growth in schizophrenia targets normative modules of synchronized development. Biol. Psychiatry 76, 438–446. doi:10.1016/j.biopsych.2014.02.010

Appelbaum, M.I., McCall, R.B., 1983. Design and analysis in developmental psychology., in: Mussen, P.H. (Ed.), Handbook of Childpsychology. Wiley, New York, pp. 415–476.

Atkinson, D., Hill, D.L.G., Stoyle, P.N.R., Summers, P.E., Clare, S., Bowtell, R., Keevil, S.F., 1999. Automatic compensation of motion artifacts in MRI. Magn. Reson. Med. 41, 163–170. doi:10.1002/(SICI)1522-2594(199901)41:1<163::AID-MRM23>3.0.CO;2- 9

Aubert-Broche, B., Fonov, V.S., García-Lorenzo, D., Mouiha, A., Guizard, N., Coupé, P., Eskildsen, S.F., Collins, D.L., 2013. A new method for structural volume analysis of longitudinal brain MRI data and its application in studying the growth trajectories of anatomical brain structures in childhood. NeuroImage 82, 393–402. doi:10.1016/j.neuroimage.2013.05.065

Backhausen, L.L., Herting, M.M., Buse, J., Roessner, V., Smolka, M.N., Vetter, N.C., 2016. Quality Control of Structural MRI Images Applied Using FreeSurfer—A Hands-On Workflow to Rate Motion Artifacts. Front. Neurosci. 10. doi:10.3389/fnins.2016.00558

Bell, R.Q., 1954. An experimental test of the accelerated longitudinal approach. Child Dev. 25, 281–286.

Bernstein, M.A., Huston, J., Ward, H.A., 2006. Imaging artifacts at 3.0T. J. Magn. Reson. Imaging JMRI 24, 735–746. doi: 10.1002/jmri.20698

Blumenthal, J.D., Zijdenbos, A., Molloy, E., Giedd, J.N., 2002. Motion artifact in magnetic resonance imaging: implications for automated analysis. NeuroImage 16, 89–92. doi:10.1006/nimg.2002.1076

Bordens, K., Abbott, B.B., 2013. Research Design and Methods: A Process Approach, 9 edition. ed. McGraw-Hill Education, New York, NY.

Braams, B.R., Duijvenvoorde, A.C.K. van, Peper, J.S., Crone, E.A., 2015. Longitudinal Changes in Adolescent Risk-Taking: A Comprehensive Study of Neural Responses to Rewards, Pubertal Development, and Risk-Taking Behavior. J. Neurosci. 35, 7226–7238. doi:10.1523/JNEUROSCI.4764-14.2015

Burgoyne, R.D., Graham, M.E., Cambray-Deakin, M., 1993. Neurotrophic effects of NMDA receptor activation on developing cerebellar granule cells. J. Neurocytol. 22, 689–695. doi:10.1 007/BF0 1181314

Button, K.S., Ioannidis, J.P.A., Mokrysz, C., Nosek, B.A., Flint, J., Robinson, E.S.J., Munafó, M.R., 2013. Power failure: why small sample size undermines the reliability of neuroscience. Nat. Rev. Neurosci. 14, 365–376. doi:10.1038/nrn3475

Cao, B., Mwangi, B., Hasan, K.M., Selvaraj, S., Zeni, C.P., Zunta-Soares, G.B., Soares, J.C., 2015. Development and validation of a brain maturation index using longitudinal neuroanatomical scans. NeuroImage 117, 311–318. doi: 10.1016/j.neuroimage.2015.05.071

Chen, Z.J., He, Y., Rosa-Neto, P., Gong, G., Evans, A.C., 2011. Age-related alterations in the modular organization of structural cortical network by using cortical thickness from MRI. NeuroImage 56, 235–245. doi: 10.1016/j.neuroimage.2011.01.010

Courchesne, E., Chisum, H.J., Townsend, J., Cowles, A., Covington, J., Egaas, B., Harwood, M., Hinds, S., Press, G.A., 2000. Normal brain development and aging: quantitative analysis at in vivo MR imaging in healthy volunteers. Radiology 216, 672–682. doi:10.1148/radiology.216.3.r00au37672

Dehaene, S., Pegado, F., Braga, L.W., Ventura, P., Filho, G.N., Jobert, A., Dehaene-Lambertz, G., Kolinsky, R., Morais, J., Cohen, L., 2010. How Learning to Read Changes the Cortical Networks for Vision and Language. Science 330, 1359–1364. doi:10.1126/science.1194140

Dennison, M., Whittle, S., Yücel, M., Vijayakumar, N., Kline, A., Simmons, J., Allen, N.B., 2013. Mapping subcortical brain maturation during adolescence: evidence of hemisphere- and sex-specific longitudinal changes. Dev. Sci. 16, 772–791. doi:10.1111/desc.12057

Diedrichsen, J., 2006. A spatially unbiased atlas template of the human cerebellum. NeuroImage 33, 127–138. doi:10.1016/j.neuroimage.2006.05.056

Draganski, B., Gaser, C., Busch, V., Schuierer, G., Bogdahn, U., May, A., 2004. Neuroplasticity: Changes in grey matter induced by training. Nature 427, 311–312. doi: 10.1038/427311a

Ducharme, S., Albaugh, M.D., Hudziak, J.J., Botteron, K.N., Nguyen, T.-V., Truong, C., Evans, A.C., Karama, S., Brain Development Cooperative Group, 2014. Anxious/depressed symptoms are linked to right ventromedial prefrontal cortical thickness maturation in healthy children and young adults. Cereb. Cortex N. Y. N 1991 24, 2941–2950. doi:10.1093/cercor/bht151

Ducharme, S., Albaugh, M.D., Nguyen, T.-V., Hudziak, J.J., Mateos-Pérez, J.M., Labbe, A., Evans, A.C., Karama, S., 2016. Trajectories of cortical thickness maturation in normal brain development — The importance of quality control procedures. NeuroImage 125, 267–279. doi: 10.1016/j.neuroimage.2015. 10.0 10

Ducharme, S., Hudziak, J.J., Botteron, K.N., Albaugh, M.D., Nguyen, T.-V., Karama, S., Evans, A.C., Brain Development Cooperative Group, 2012. Decreased regional cortical thickness and thinning rate are associated with inattention symptoms in healthy children. J. Am. Acad. Child Adolesc. Psychiatry 51, 18–27.e2. doi: 10.1016/j jaac.2011.09.022

Evans, A.C., 2013. Networks of anatomical covariance. NeuroImage 80, 489–504. doi: 10.1016/j.neuroimage.2013.05.054

Finlay, B.L., Darlington, R.B., 1995. Linked regularities in the development and evolution of mammalian brains. Science 268, 1578–1584.

Fjell, A.M., Grydeland, H., Krogsrud, S.K., Amlien, I., Rohani, D.A., Ferschmann, L., Storsve, A.B., Tamnes, C.K., Sala-Llonch, R., Due-Tønnessen, P., Bjørnerud, A., Sølsnes, A.E., Háberg, A.K., Skranes, J., Bartsch, H., Chen, C.-H., Thompson, W.K., Panizzon, M.S., Kremen, W.S., Dale, A.M., Walhovd, K.B., 2015. Development and aging of cortical thickness correspond to genetic organization patterns. Proc. Natl. Acad. Sci. U. S. A. 112, 15462–15467. doi: 10.1073/pnas.1508831112

Friedel, S., Whittle, S.L., Vijayakumar, N., Simmons, J.G., Byrne, M.L., Schwartz, O.S., Allen, N.B., 2015. Dispositional mindfulness is predicted by structural development of the insula during late adolescence. Dev. Cogn. Neurosci. 14, 62–70. doi:10.1016/j.dcn.2015.07.001

Galbraith, S., Bowden, J., Mander, A., 2017. Accelerated longitudinal designs: An overview of modelling, power, costs and handling missing data. Stat. Methods Med. Res. 26, 374–398. doi:10.1177/0962280214547150

Gaser, C., Schlaug, G., 2003. Brain Structures Differ between Musicians and Non-Musicians. J. Neurosci. 23, 9240–9245.

Geng, X., Li, G., Lu, Z., Gao, W., Wang, L., Shen, D., Zhu, H., Gilmore, J.H., 2016. Structural and Maturational Covariance in Early Childhood Brain Development. Cereb. Cortex N. Y. N 1991. doi: 10.1093/cercor/bhw022

Gibbons, R.D., Hedeker, D., DuToit, S., 2010. Advances in Analysis of Longitudinal Data. Annu. Rev. Clin. Psychol. 6, 79–107. doi: 10.1146/annurev.clinpsy.032408.153550

Giedd, J.N., Blumenthal, J., Jeffries, N.O., Castellanos, F.X., Liu, H., Zijdenbos, A., Paus, T., Evans, A.C., Rapoport, J.L., 1999. Brain development during childhood and adolescence: a longitudinal MRI study. Nat. Neurosci. 2, 861–863. doi:10.1038/13158

Giedd, J.N., Raznahan, A., Mills, K.L., Lenroot, R.K., 2012. Review: magnetic resonance imaging of male/female differences in human adolescent brain anatomy. Biol. Sex Differ. 3, 19. doi:10.1186/2042-6410-3-19

Goddings, A.-L., Mills, K.L., Clasen, L.S., Giedd, J.N., Viner, R.M., Blakemore, S.-J., 2014. The influence of puberty on subcortical brain development. NeuroImage 88, 242–251. doi: 10.1016/j.neuroimage.2013.09.073

Gogtay, N., Giedd, J.N., Lusk, L., Hayashi, K.M., Greenstein, D., Vaituzis, A.C., Nugent, T.F., Herman, D.H., Clasen, L.S., Toga, A.W., Rapoport, J.L., Thompson, P.M., 2004. Dynamic mapping of human cortical development during childhood through early adulthood. Proc. Natl. Acad. Sci. U. S. A. 101, 8174–8179. doi:10.1073/pnas.0402680101

Gogtay, N., Nugent, T.F., Herman, D.H., Ordonez, A., Greenstein, D., Hayashi, K.M., Clasen, L., Toga, A.W., Giedd, J.N., Rapoport, J.L., Thompson, P.M., 2006. Dynamic mapping of normal human hippocampal development. Hippocampus 16, 664–672. doi:10.1002/hipo.20193

Gong, G., He, Y., Chen, Z.J., Evans, A.C., 2012. Convergence and divergence of thickness correlations with diffusion connections across the human cerebral cortex. NeuroImage 59, 1239–1248. doi:10.1016/j.neuroimage.2011.08.017

Gorgolewski, K.J., Alfaro-Almagro, F., Auer, T., Bellec, P., Capotă, M., Chakravarty, M.M., Churchill, N.W., Cohen, A.L., Craddock, R.C., Devenyi, G.A., Eklund, A., Esteban, O., Flandin, G., Ghosh, S.S., Guntupalli, J.S., Jenkinson, M., Keshavan, A., Kiar, G., Liem, F., Raamana, P.R., Raffelt, D., Steele, C.J., Quirion, P.-O., Smith, R.E., Strother, S.C., Varoquaux, G., Wang, Y., Yarkoni, T., Poldrack, R.A., 2017. BIDS apps: Improving ease of use, accessibility, and reproducibility of neuroimaging data analysis methods. PLOS Comput. Biol. 13, e1005209. doi:10.1371 /journal.pcbi.1005209

Greene, D.J., Black, K.J., Schlaggar, B.L., 2016. Considerations for MRI study design and implementation in pediatric and clinical populations. Dev. Cogn. Neurosci. 18, 101–112. doi:10.1016/j.dcn.2015.12.005

Greenstein, D., Lerch, J., Shaw, P., Clasen, L., Giedd, J., Gochman, P., Rapoport, J., Gogtay, N., 2006. Childhood onset schizophrenia: cortical brain abnormalities as young adults. J. Child Psychol. Psychiatry 47, 1003–1012. doi:10.1111/j.1469-7610.2006.01658.x

Han, X., Jovicich, J., Salat, D., van der Kouwe, A., Quinn, B., Czanner, S., Busa, E., Pacheco, J., Albert, M., Killiany, R., Maguire, P., Rosas, D., Makris, N., Dale, A., Dickerson, B., Fischl, B., 2006. Reliability of MRI-derived measurements of human cerebral cortical thickness: the effects of field strength, scanner upgrade and manufacturer. NeuroImage 32, 180–194. doi: 10.1016/j.neuroimage.2006.02.051

Hardan, A.Y., Muddasani, S., Vemulapalli, M., Keshavan, M.S., Minshew, N.J., 2006. An MRI Study of Increased Cortical Thickness in Autism. Am. J. Psychiatry 163, 1290–1292. doi: 10.1 176/appi.ajp.163.7.1290

Heinen, R., Bouvy, W.H., Mendrik, A.M., Viergever, M.A., Biessels, G.J., Bresser, J. de, 2016. Robustness of Automated Methods for Brain Volume Measurements across Different MRI Field Strengths. PLOS ONE 11, e0165719. doi:10.1371 /journal.pone.0165719

Herting, M.M., Gautam, P., Spielberg, J.M., Dahl, R.E., Sowell, E.R., 2015. A Longitudinal Study: Changes in Cortical Thickness and Surface Area during Pubertal Maturation. PLOS ONE 10, e0119774. doi: 10.1371/journal.pone.01 19774

Herting, M.M., Gautam, P., Spielberg, J.M., Kan, E., Dahl, R.E., Sowell, E.R., 2014. The role of testosterone and estradiol in brain volume changes across adolescence: a longitudinal structural MRI study. Hum. Brain Mapp. 35, 5633–5645. doi: 10.1002/hbm.22575

Iglesias, J.E., Augustinack, J.C., Nguyen, K., Player, C.M., Player, A., Wright, M., Roy, N., Frosch, M.P., McKee, A.C., Wald, L.L., Fischl, B., Van Leemput, K., Alzheimer’s Disease Neuroimaging Initiative, 2015. A computational atlas of the hippocampal formation using ex vivo, ultra-high resolution MRI: Application to adaptive segmentation of in vivo MRI. NeuroImage 115, 117–137. doi: 10.1016/j.neuroimage.2015.04.042

Im, K., Lee, J.-M., Lyttelton, O., Kim, S.H., Evans, A.C., Kim, S.I., 2008. Brain Size and Cortical Structure in the Adult Human Brain. Cereb. Cortex 18, 2181–2191. doi: 10.1093/cercor/bhm244

Jahanshad, N., Thompson, P.M., 2017. Multimodal neuroimaging of male and female brain structure in health and disease across the life span. J. Neurosci. Res. 95, 371–379. doi: 10.1002/jnr.23919

Jovicich, J., Czanner, S., Han, X., Salat, D., van der Kouwe, A., Quinn, B., Pacheco, J., Albert, M., Killiany, R., Blacker, D., Maguire, P., Rosas, D., Makris, N., Gollub, R., Dale, A., Dickerson, B.C., Fischl, B., 2009. MRI-derived measurements of human subcortical, ventricular and intracranial brain volumes: Reliability effects of scan sessions, acquisition sequences, data analyses, scanner upgrade, scanner vendors and field strengths. NeuroImage 46, 177–192. doi: 10.1016/j.neuroimage.2009.02.010

Jovicich, J., Marizzoni, M., Sala-Llonch, R., Bosch, B., Bartrés-Faz, D., Arnold, J., Benninghoff, J., Wiltfang, J., Roccatagliata, L., Nobili, F., Hensch, T., Tränkner, A., Schönknecht, P., Leroy, M., Lopes, R., Bordet, R., Chanoine, V., Ranjeva, J.-P., Didic, M., Gros-Dagnac, H., Payoux, P., Zoccatelli, G., Alessandrini, F., Beltramello, A., Bargall—, N., Blin, O., Frisoni, G.B., PharmaCog Consortium, 2013. Brain morphometry reproducibility in multi-center 3T MRI studies: a comparison of cross-sectional and longitudinal segmentations. NeuroImage 83, 472–484. doi: 10.1016/j.neuroimage.2013.05.007

Kelly, C., Toro, R., Di Martino, A., Cox, C.L., Bellec, P., Castellanos, F.X., Milham, M.P., 2012. A convergent functional architecture of the insula emerges across imaging modalities. NeuroImage 61, 1129–1142. doi:10.1016/j.neuroimage.2012.03.021

Khundrakpam, B.S., Reid, A., Brauer, J., Carbonell, F., Lewis, J., Ameis, S., Karama, S., Lee, J., Chen, Z., Das, S., Evans, A.C., Brain Development Cooperative Group, 2013. Developmental changes in organization of structural brain networks. Cereb. Cortex N. Y. N 1991 23, 2072–2085. doi: 10.1093/cercor/bhs 187

Korin, H.W., Felmlee, J.P., Ehman, R.L., Riederer, S.J., 1990. Adaptive technique for three-dimensional MR imaging of moving structures. Radiology 177, 217–221. doi: 10.1148/radiology.177.1.2399320

Kraemer, H.C., Yesavage, J.A., Taylor, J.L., Kupfer, D., 2000. How Can We Learn About Developmental Processes From Cross-Sectional Studies, or Can We? Am. J. Psychiatry 157, 163–171. doi: 10.1176/appi.ajp.157.2.163

Krongold, M., Cooper, C., Bray, S., 2015. Modular Development of Cortical Gray Matter Across Childhood and Adolescence. Cereb. Cortex bhv 307. doi:10.1093/cercor/bhv3 07

Kruggel, F., Turner, J., Muftuler, L.T., Alzheimer’s Disease Neuroimaging Initiative, 2010. Impact of scanner hardware and imaging protocol on image quality and compartment volume precision in the ADNI cohort. NeuroImage 49, 2123–2133. doi: 10.1016/j.neuroimage.2009.11.006

Lebel, C., Beaulieu, C., 2011. Longitudinal development of human brain wiring continues from childhood into adulthood. J. Neurosci. Off. J. Soc. Neurosci. 31, 10937–10947. doi: 10.1 523/JNEUROSCI.5302-10.2011

Lenroot, R.K., Gogtay, N., Greenstein, D.K., Wells, E.M., Wallace, G.L., Clasen, L.S., Blumenthal, J.D., Lerch, J., Zijdenbos, A.P., Evans, A.C., Thompson, P.M., Giedd, J.N., 2007. Sexual dimorphism of brain developmental trajectories during childhood and adolescence. NeuroImage 36, 1065–1073. doi:10.1016/j.neuroimage.2007.03.053

Leonard, C.M., Towler, S., Welcome, S., Halderman, L.K., Otto, R., Eckert, M.A., Chiarello, C., 2008. Size Matters: Cerebral Volume Influences Sex Differences in Neuroanatomy. Cereb. Cortex N. Y. NY 18, 2920–2931. doi:10.1093/cercor/bhn052

Li, X., Pu, F., Fan, Y., Niu, H., Li, S., Li, D., 2013. Age-related changes in brain structural covariance networks. Front. Hum. Neurosci. 7. doi:10.3389/fnhum.2013.00098

Madhyastha T, Peverill M, Koh N, McCabe C, Flournoy JC, Mills KL, King K, & McLaughlin K. (in revision). Current methods and limitations for longitudinal fMRI analysis across development. Developmental Cognitive Neuroscience.

Marshall, W.A., Tanner, J.M., 1969. Variations in pattern of pubertal changes in girls. Arch. Dis. Child. 44, 291–303.

Mills, K.L., Goddings, A.-L., Herting, M.M., Meuwese, R., Blakemore, S.-J., Crone, E.A., Dahl, R.E., Güroğlu, B., Raznahan, A., Sowell, E.R., Tamnes, C.K., 2016. Structural brain development between childhood and adulthood: Convergence across four longitudinal samples. Neuroimage 141, 273–281. doi: 10.1016/j.neuroimage.2016.07.044

Mills, K.L., Goddings, A.-L., Clasen, L.S., Giedd, J.N., Blakemore, S.-J., 2014a. The Developmental Mismatch in Structural Brain Maturation during Adolescence. Dev. Neurosci. 36, 147–160. doi: 10.1159/000362328

Mills, K.L., Lalonde, F., Clasen, L.S., Giedd, J.N., Blakemore, S.-J., 2014b. Developmental changes in the structure of the social brain in late childhood and adolescence. Soc. Cogn. Affect. Neurosci. 9, 123–131. doi: 10.1093/scan/nss113

Mills, K.L., Tamnes, C.K., 2014. Methods and considerations for longitudinal structural brain imaging analysis across development. Dev. Cogn. Neurosci. 9, 172–190. doi: 10.1016/j.dcn.2014.04.004

Morey, R.A., Selgrade, E.S., Wagner, H.R., Huettel, S.A., Wang, L., McCarthy, G., 2010. Scan-rescan reliability of subcortical brain volumes derived from automated segmentation. Hum. Brain Mapp. 31, 1751–1762. doi: 10.1002/hbm.20973

Mutlu, A.K., Schneider, M., Debbané, M., Badoud, D., Eliez, S., Schaer, M., 2013. Sex differences in thickness, and folding developments throughout the cortex. NeuroImage 82, 200–207. doi: 10.1016/j.neuroimage.2013.05.076

Nie, J., Li, G., Shen, D., 2013. Development of cortical anatomical properties from early childhood to early adulthood. NeuroImage 76, 216–224. doi:10.1016/j.neuroimage.2013.03.021

Nguyen, T.-V., McCracken, J., Ducharme, S., Botteron, K.N., Mahabir, M., Johnson, W., Israel, M., Evans, A.C., Karama, S., Group, for the B.D.C., 2013a. Testosterone-Related Cortical Maturation Across Childhood and Adolescence. Cereb. Cortex 23, 1424–1432. doi:10.1093/cercor/bhs125

Nguyen, T.-V., McCracken, J.T., Ducharme, S., Cropp, B.F., Botteron, K.N., Evans, A.C., Karama, S., 2013b. Interactive Effects of Dehydroepiandrosterone and Testosterone on Cortical Thickness during Early Brain Development. J. Neurosci. 33, 10840– 10848. doi:10.1523/JNEUROSCI.5747-12.2013

Ordaz, S.J., Foran, W., Velanova, K., Luna, B., 2013. Longitudinal Growth Curves of Brain Function Underlying Inhibitory Control through Adolescence. J. Neurosci. 33, 18109—18124. doi: 10.1523/JNEUROSCI.1741-13.2013

Palaniyappan, L., 2010. Computing cortical surface measures in schizophrenia. Br. J. Psychiatry 196, 414–414. doi:10.1192/bjp.196.5.414

Pardoe, H.R., Kucharsky Hiess, R., Kuzniecky, R., 2016. Motion and morphometry in clinical and nonclinical populations. NeuroImage 135, 177–185. doi:10.1016/j.neuroimage.2016.05.005

Paus, T., Wong, A.P.-Y., Syme, C., Pausova, Z., 2017. Sex differences in the adolescent brain and body: Findings from the saguenay youth study. J. Neurosci. Res. 95, 362–370. doi: 10.1002/jnr.23825

Pfefferbaum, A., Mathalon, D.H., Sullivan, E.V., Rawles, J.M., Zipursky, R.B., Lim, K.O., 1994. A quantitative magnetic resonance imaging study of changes in brain morphology from infancy to late adulthood. Arch. Neurol. 51, 874–887.

Pinheiro, J., Bates, D., 2013. Mixed-Effects Models in S and S-PLUS, Softcover reprint of the original 1st ed. 2000 edition. ed. Springer, New York, NY u.a.

Power, J.D., Barnes, K.A., Snyder, A.Z., Schlaggar, B.L., Petersen, S.E., 2013. Steps toward optimizing motion artifact removal in functional connectivity MRI; a reply to Carp. NeuroImage 76. doi:10.1016/j.neuroimage.2012.03.017

Power, J.D., Barnes, K.A., Snyder, A.Z., Schlaggar, B.L., Petersen, S.E., 2012. Spurious but systematic correlations in functional connectivity MRI networks arise from subject motion. NeuroImage 59, 2142–2154. doi:10.1016/j.neuroimage.2011.10.018

Qin, L., van Gelderen, P., Derbyshire, J.A., Jin, F., Lee, J., de Zwart, J.A., Tao, Y., Duyn, J.H., 2009. Prospective head-movement correction for high-resolution MRI using an in-bore optical tracking system. Magn. Reson. Med. 62, 924–934. doi:10.1002/mrm.22076

Rakic, P., 1988. Specification of cerebral cortical areas. Science 241, 170–176.

Raschle, N., Zuk, J., Ortiz-Mantilla, S., Sliva, D.D., Franceschi, A., Grant, P.E., Benasich, A.A., Gaab, N., 2012. Pediatric neuroimaging in early childhood and infancy: challenges and practical guidelines. Ann. N. Y. Acad. Sci. 1252, 43–50. doi:10.1111/j.1749-6632.2012.06457.x

Raznahan, A., Lerch, J.P., Lee, N., Greenstein, D., Wallace, G.L., Stockman, M., Clasen, L., Shaw, P.W., Giedd, J.N., 2011a. Patterns of Coordinated Anatomical Change in Human Cortical Development: A Longitudinal Neuroimaging Study of Maturational Coupling. Neuron 72, 873–884. doi: 10.1016/j.neuron.2011.09.028

Raznahan, A., Shaw, P., Lalonde, F., Stockman, M., Wallace, G.L., Greenstein, D., Clasen, L., Gogtay, N., Giedd, J.N., 2011b. How Does Your Cortex Grow? J. Neurosci. 31, 7174–7177. doi:10.1523/JNEUROSCI.0054-11.2011

Raznahan, A., Shaw, P.W., Lerch, J.P., Clasen, L.S., Greenstein, D., Berman, R., Pipitone, J., Chakravarty, M.M., Giedd, J.N., 2014. Longitudinal four-dimensional mapping of subcortical anatomy in human development. Proc. Natl. Acad. Sci. 111, 1592–1597. doi: 10.1073/pnas.1316911111

Reardon, P.K., Clasen, L., Giedd, J.N., Blumenthal, J., Lerch, J.P., Chakravarty, M.M., Raznahan, A., 2016. An Allometric Analysis of Sex and Sex Chromosome Dosage Effects on Subcortical Anatomy in Humans. J. Neurosci. Off. J. Soc. Neurosci. 36, 2438–2448. doi:10.1523/JNEUROSCI.3195-15.2016

Reiss, P.T., Huang, L., Chen, Y.-H., Huo, L., Tarpey, T., Mennes, M., 2014. Massively parallel nonparametric regression, with an application to developmental brain mapping. J. Comput. Graph. Stat. Jt. Publ. Am. Stat. Assoc. Inst. Math. Stat. Interface Found. N. Am. 23, 232–248. doi:10.1080/10618600.2012.733549

Reuter, M., Fischl, B., 2011. Avoiding asymmetry-induced bias in longitudinal image processing. NeuroImage 57, 19–21. doi:10.1016/j.neuroimage.2011.02.076

Reuter, M., Schmansky, N.J., Rosas, H.D., Fischl, B., 2012. Within-subject template estimation for unbiased longitudinal image analysis. Neuroimage 61, 1402–1418. doi: 10.1016/j.neuroimage.2012.02.084

Reuter, M., Tisdall, M.D., Qureshi, A., Buckner, R.L., van der Kouwe, A.J.W., Fischl, B., 2015. Head motion during MRI acquisition reduces gray matter volume and thickness estimates. NeuroImage 107, 107–115. doi:10.1016/j.neuroimage.2014.12.006

Riska, B., 1986. Some Models for Development, Growth, and Morphometric Correlation. Evolution 40, 1303–1311. doi:10.2307/2408955

Sanabria-Diaz, G., Melie-Garcìa, L., Iturria-Medina, Y., Alemán-G—mez, Y., Hernández-González, G., Valdés-Urrutia, L., Galán, L., Valdés-Sosa, P., 2010. Surface area and cortical thickness descriptors reveal different attributes of the structural human brain networks. NeuroImage 50, 1497–1510. doi:10.1016/j.neuroimage.2010.01.028

Satterthwaite, T.D., Elliott, M.A., Gerraty, R.T., Ruparel, K., Loughead, J., Calkins, M.E., Eickhoff, S.B., Hakonarson, H., Gur, R.C., Gur, R.E., Wolf, D.H., 2013. An Improved Framework for Confound Regression and Filtering for Control of Motion Artifact in the Preprocessing of Resting-State Functional Connectivity Data. NeuroImage 64. doi: 10. 10 1 6/j.neuroimage.2012.08.052

Savalia, N.K., Agres, P.F., Chan, M.Y., Feczko, E.J., Kennedy, K.M., Wig, G.S., 2017. Motion-related artifacts in structural brain images revealed with independent estimates of in-scanner head motion. Hum. Brain Mapp. 38, 472–492. doi:10.1002/hbm.23397

Seeley, W.W., Crawford, R.K., Zhou, J., Miller, B.L., Greicius, M.D., 2009. Neurodegenerative diseases target large-scale human brain networks. Neuron 62, 42–52. doi: 10.1016/j.neuron.2009.03.024

Shaw, P., Eckstrand, K., Sharp, W., Blumenthal, J., Lerch, J.P., Greenstein, D., Clasen, L., Evans, A., Giedd, J., Rapoport, J.L., 2007. Attention-deficit/hyperactivity disorder is characterized by a delay in cortical maturation. Proc. Natl. Acad. Sci. 104, 19649–19654. doi:10.1073/pnas.0707741104

Shaw, P., Gilliam, M., Liverpool, M., Weddle, C., Malek, M., Sharp, W., Greenstein, D., Evans, A., Rapoport, J., Giedd, J., 2011. Cortical Development in Typically Developing Children With Symptoms of Hyperactivity and Impulsivity: Support for a Dimensional View of Attention Deficit Hyperactivity Disorder. Am. J. Psychiatry 168, 143–151. doi:10.1176/appi.ajp.2010.10030385

Shaw, P., Greenstein, D., Lerch, J., Clasen, L., Lenroot, R., Gogtay, N., Evans, A., Rapoport, J., Giedd, J., 2006. Intellectual ability and cortical development in children and adolescents. Nature 440, 676–679. doi:10.1038/nature04513

Shaw, P., Kabani, N.J., Lerch, J.P., Eckstrand, K., Lenroot, R., Gogtay, N., Greenstein, D., Clasen, L., Evans, A., Rapoport, J.L., Giedd, J.N., Wise, S.P., 2008. Neurodevelopmental trajectories of the human cerebral cortex. J. Neurosci. Off. J. Soc. Neurosci. 28, 3586–3594. doi:10.1523/JNEUROSCI.5309-07.2008

Shirtcliff, E.A., Dahl, R.E., Pollak, S.D., 2009. Pubertal development: correspondence between hormonal and physical development. Child Dev. 80, 327–337. doi:10.1111/j.1467-8624.2009.01263.x

Singer, J.D., Willett, J.B., 2003. Applied Longitudinal Data Analysis: Modeling Change and Event Occurrence, 1 edition. ed. Oxford; University Press, Oxford; New York.

Sowell, E.R., Thompson, P.M., Leonard, C.M., Welcome, S.E., Kan, E., Toga, A.W., 2004. Longitudinal Mapping of Cortical Thickness and Brain Growth in Normal Children. J. Neurosci. 24, 8223–8231. doi: 10.1523/JNEUROSCI. 1798-04.2004

Steen, R.G., Hamer, R.M., Lieberman, J.A., 2007. Measuring Brain Volume by MR Imaging: Impact of Measurement Precision and Natural Variation on Sample Size Requirements. Am. J. Neuroradiol. 28, 1119–1125. doi:10.3174/ajnr.A0537

Sullivan, E.V., Pfefferbaum, A., Rohlfing, T., Baker, F.C., Padilla, M.L., Colrain, I.M., 2011. Developmental change in regional brain structure over 7 months in early adolescence: Comparison of approaches for longitudinal atlas-based parcellation. NeuroImage 57, 214–224. doi:10.1016/j.neuroimage.2011.04.003

Sun, S.S., Schubert, C.M., Chumlea, W.C., Roche, A.F., Kulin, H.E., Lee, P.A., Himes, J.H., Ryan, A.S., 2002. National estimates of the timing of sexual maturation and racial differences among US children. Pediatrics 110, 911–919.

Swagerman, S.C., Brouwer, R.M., de Geus, E.J.C., Hulshoff Pol, H.E., Boomsma, D.I., 2014. Development and heritability of subcortical brain volumes at ages 9 and 12. Genes Brain Behav. 13, 733–742. doi:10.1111 /gbb.12182

Tamnes, C.K., Herting, M.M., Goddings, A.-L., Meuwese, R., Blakemore, S.-J., Dahl, R.E., Güroğlu, B., Raznahan, A., Sowell, E.R., Crone, E.A., Mills, K.L., 2017. Development of the Cerebral Cortex across Adolescence: A Multisample Study of Inter-Related Longitudinal Changes in Cortical Volume, Surface Area, and Thickness. J. Neurosci. 37, 3402–3412. doi:10.1523/JNEUROSCI.3302-16.2017

Tamnes, C.K., Walhovd, K.B., Dale, A.M., Østby, Y., Grydeland, H., Richardson, G., Westlye, L.T., Roddey, J.C., Hagler Jr., D.J., Due-Tønnessen, P., Holland, D., Fjell, A.M., 2013. Brain development and aging: Overlapping and unique patterns of change. NeuroImage 68, 63–74. doi: 10.1016/j.neuroimage.2012.11.03 9

Tanaka, C., Matsui, M., Uematsu, A., Noguchi, K., Miyawaki, T., 2012. Developmental Trajectories of the Fronto-Temporal Lobes from Infancy to Early Adulthood in Healthy Individuals. Dev. Neurosci. 34, 477–487. doi:10.1159/000345152

Thompson, P.M., Giedd, J.N., Woods, R.P., MacDonald, D., Evans, A.C., Toga, A.W., 2000. Growth patterns in the developing brain detected by using continuum mechanical tensor maps. Nature 404, 190–193. doi: 10.1038/35004593

Tiemeier, H., Lenroot, R.K., Greenstein, D.K., Tran, L., Pierson, R., Giedd, J.N., 2010. Cerebellum development during childhood and adolescence: a longitudinal morphometric MRI study. NeuroImage 49, 63–70. doi:10.1016/j.neuroimage.2009.08.016

Tijssen, R.H.N., Jansen, J.F.A., Backes, W.H., 2009. Assessing and minimizing the effects of noise and motion in clinical DTI at 3 T. Hum. Brain Mapp. 30, 2641–2655. doi: 10.1002/hbm.20695

Toro, R., Chupin, M., Garnero, L., Leonard, G., Perron, M., Pike, B., Pitiot, A., Richer, L., Veillette, S., Pausova, Z., Paus, T., 2009. Brain volumes and Val66Met polymorphism of the BDNF gene: local or global effects? Brain Struct. Funct. 213, 501–509. doi:10.1007/s00429-009-0203-y

Urosevic, S., Collins, P., Muetzel, R., Lim, K., Luciana, M., 2012. Longitudinal changes in behavioral approach system sensitivity and brain structures involved in reward processing during adolescence. Dev. Psychol. 48, 1488–1500. doi:10.1037/a0027502

Van Essen, D.C., 1997. A tension-based theory of morphogenesis and compact wiring in the central nervous system. Nature 385, 313–318. doi:10.1038/385313a0

van Soelen, I.L.C., Brouwer, R.M., van Baal, G.C.M., Schnack, H.G., Peper, J.S., Collins, D.L., Evans, A.C., Kahn, R.S., Boomsma, D.I., Hulshoff Pol, H.E., 2012. Genetic influences on thinning of the cerebral cortex during development. NeuroImage 59, 3871–3880. doi:10.1016/j.neuroimage.2011.11.044

Verbeke, G., Molenberghs, G., 2000. Linear Mixed Models for Longitudinal Data. Springer-Verlag, New York.

Vijayakumar, N., Allen, N.B., Youssef, G., Dennison, M., Yücel, M., Simmons, J.G., Whittle, S., 2016. Brain development during adolescence: A mixed-longitudinal investigation of cortical thickness, surface area, and volume. Hum. Brain Mapp. 37, 2027–2038. doi: 10.1002/hbm.23154

Vijayakumar, N., Allen, N.B., Youssef, G.J., Simmons, J.G., Byrne, M.L., Whittle, S., 2016. Neurodevelopmental Trajectories Related to Attention Problems Predict Driving-Related Risk Behaviors. J. Atten. Disord. 1087054716682336. doi:10.1177/ 1087054716682336

Vijayakumar, N., Whittle, S., Dennison, M., Yücel, M., Simmons, J., Allen, N.B., 2014a. Development of temperamental effortful control mediates the relationship between maturation of the prefrontal cortex and psychopathology during adolescence: A 4- year longitudinal study. Dev. Cogn. Neurosci. 9, 30–43. doi:10.1016/j.dcn.2013.12.002

Vijayakumar, N., Whittle, S., Yücel, M., Dennison, M., Simmons, J., Allen, N.B., 2014b.Thinning of the lateral prefrontal cortex during adolescence predicts emotion regulation in females. Soc. Cogn. Affect. Neurosci. 9, 1845–1854. doi:10.1093/scan/nst183

Walhovd, K.B., Fjell, A.M., Giedd, J., Dale, A.M., Brown, T.T., 2016. Through thick and thin: a need to reconcile contradictory results on trajectories in human cortical development. Cereb. Cortex bhv 301.

Walhovd, K.B., Tamnes, C.K., Bjørnerud, A., Due-Tønnessen, P., Holland, D., Dale, A.M., Fjell, A.M., 2015. Maturation of Cortico-Subcortical Structural Networks— Segregation and Overlap of Medial Temporal and Fronto-Striatal Systems in Development. Cereb. Cortex 25, 1835–1841. doi: 10.1093/cercor/bht424

Wallace, G.L., Dankner, N., Kenworthy, L., Giedd, J.N., Martin, A., 2010. Age-related temporal and parietal cortical thinning in autism spectrum disorders. Brain J. Neurol. 133, 3745–3754. doi:10.1093/brain/awq279

West, B.T., Welch, K.B., Galecki, A.T., 2006. Linear Mixed Models: A Practical Guide Using Statistical Software, 1 edition. ed. Chapman and Hall/CRC, Boca Raton.

Westbrook, C., Roth, C.K., Talbot, J., 2011. MRI in practice, 4th ed. Blackwell Publishing, Oxford.

White, N., Roddey, C., Shankaranarayanan, A., Han, E., Rettmann, D., Santos, J., Kuperman, J., Dale, A., 2010. PROMO: Real-time prospective motion correction in MRI using image-based tracking. Magn. Reson. Med. 63, 91–105. doi:10.1002/mrm.22176

Wierenga, L.M., Langen, M., Ambrosino, S., van Dijk, S., Oranje, B., Durston, S., 2014a. Typical development of basal ganglia, hippocampus, amygdala and cerebellum from age 7 to 24. NeuroImage 96, 67–72. doi:10.1016/j.neuroimage.2014.03.072

Wierenga, L.M., Langen, M., Oranje, B., Durston, S., 2014b. Unique developmental trajectories of cortical thickness and surface area. NeuroImage 87, 120–126. doi:10.1016/j.neuroimage.2013.11.010

Wonderlick, J.S., Ziegler, D.A., Hosseini-Varnamkhasti, P., Locascio, J.J., Bakkour, A., van der Kouwe, A., Triantafyllou, C., Corkin, S., Dickerson, B.C., 2009. Reliability of MRI-derived cortical and subcortical morphometric measures: effects of pulse sequence, voxel geometry, and parallel imaging. NeuroImage 44, 1324–1333. doi:10.1016/j.neuroimage.2008.10.03 7

Zhang, K., Sejnowski, T.J., 2000. A universal scaling law between gray matter and white matter of cerebral cortex. Proc. Natl. Acad. Sci. U. S. A. 97, 5621–5626.

Zhou, D., Lebel, C., Treit, S., Evans, A., Beaulieu, C., 2015. Accelerated longitudinal cortical thinning in adolescence. NeuroImage 104, 138–145. doi:10.1016/j.neuroimage.2014.10.005

Ziegler, G., Ridgway, G., Blakemore, S.J., Ashburner, J., Penny, W., 2016. Multivariate dynamical modelling of structural change during development 147, 746–762–746– 762.

Zielinski, B.A., Gennatas, E.D., Zhou, J., Seeley, W.W., 2010. Network-level structural covariance in the developing brain. Proc. Natl. Acad. Sci. U. S. A. 107, 18191–18196. doi:10.1073/pnas.1003109107

